# Transplanted human photoreceptors differentially survive, incorporate, and mature in mildly and severely degenerated mouse retinae

**DOI:** 10.64898/2026.06.18.733059

**Authors:** Maria Pavlou, Karen Teßmer, Juliane Hammer, Thomas Kurth, Aikaterini Makri, Celine Palitza, Berta Coll San Martin, Fabian Rost, Marius Ader

## Abstract

Photoreceptor transplantation is considered a disease-agnostic therapeutic strategy for retinal degenerative diseases with highly heterogenous genetic, molecular, and cellular pathologies. While integration of human photoreceptors enriched from stem cell-derived retinal organoids was noted in previous preclinical studies, the potential influence of retinal degeneration severity on transplantation efficiency has not been systematically assessed. Here, we employed mice presenting mild or severe retinal degeneration as recipients for human induced pluripotent stem cell-derived photoreceptors. Donor cells formed multi-cellular clusters that structurally integrated from 3 weeks post-transplantation (wpt) in mildly degenerated retinas, closely interacting with host Müller glia, resulting in proper maturation characterized by inner/outer segment and synapse formation by 26 wpt. In contrast, in severely degenerated hosts, donor photoreceptors remained mainly singularized and scattered in the subretinal space, showing limited structural integration or signs of maturation. Differential maturation of donor cells in mild vs. severe hosts was confirmed by single-cell RNA-sequencing analysis. However, transplantation at the beginning of the degeneration process of the severe model allowed structural integration and maturation of donor photoreceptors, despite complete loss of endogenous photoreceptors over time. The study thus shows that survival, integration, and maturation of donor photoreceptors depend on the degenerative retinal microenvironment shaping significantly transplantation efficiency.

## INTRODUCTION

Retinal degenerative diseases (RDDs), such as retinitis pigmentosa and age-related macular degeneration, culminate in loss of photoreceptors, the neuronal light-sensitive cells of the retina. As this cell type cannot regenerate in the mammalian retina, cell replacement is considered as a general disease-agnostic therapeutic strategy for vision restoration. Emphasis has so far been placed on the generation of candidate donor cells from embryonic (ESC) or induced pluripotent stem cells (iPSC), in order to produce retinal organoids and isolate a transplantable population of photoreceptors. Indeed, previous studies provide evidence for incorporation of human ESC-/iPSC-derived photoreceptors into mouse models of retinal degeneration and restoration of some photoreceptor-mediated vision,^1–4^ culminating in the initiation of a recently started first clinical trial (NCT06789445, BlueRock Therapeutics, ClinicalTrials.gov).

RDD pathologies are characterized by high causative and phenotypic heterogeneity and thus diverse extent of photoreceptor degeneration, accompanied by inner retinal remodelling. Recent transplantation studies used different RDD mouse models ranging from moderate to severe photoreceptor loss.^5^ While signs of donor photoreceptor incorporation including synapse formation, inner-/outer-segment generation, and interaction with Müller glia cells as well as some functional rescue were reported, signs of photoreceptor maturation and the degree of donor cell integration appeared highly various in the different studies.^2,4,6,7^ Thus, there is currently limited knowledge to what extend the degeneration status influences donor-host interaction and maturation of donor photoreceptors, parameters which are key for functional integration and vision restoration.

To systematically evaluate the influence of the degeneration status on photoreceptor transplants, we utilised in this study the cone-deficient mouse model Cone photoreceptor function loss 1 (Cpfl1)^8^, and the cone- and rod-deficient mouse model Cpfl1/Rhodopsin^-/-^ (tg(Cpfl1/Rho^-/-^)),^9^ representing mild and severe retinal degeneration, respectively. Donor photoreceptors were isolated from retinal organoids differentiated from a human photoreceptor-specific reporter iPSC line (CRX-mCherry-iPSC),^10,11^ followed by subretinal delivery into host mice.^2,12^ Histological analysis and single-cell gene expression profiling revealed that transplanted photoreceptors can survive long-term (up to six months) in both mildly and severely degenerated retinae, however, significantly differ in their distribution, integration, and differentiation status, with drastically improved cluster formation, donor-host interaction, and photoreceptor maturation within the mild degeneration environment of Cpfl1 mice. However, when transplanted at an early stage of retinal degeneration, cluster formation, structural integration, and maturation of donor cells was also observed in the severe tg(Cpfl1/Rho^-/-^) model, despite complete loss of endogenous photoreceptors over time.

## RESULTS

### Validation of a human photoreceptor reporter iPSC line to produce a transplantable population of photoreceptors

A human iPSC line, bearing the mCherry tag under the control of a photoreceptor-specific cone-rod homeobox (CRX) promoter was used to produce human retinal organoids (HROs).^10^ Day (D) 200 HROs present a laminated structure, resembling organisation of the human retina, where Müller glia nuclei (SOX2) reside beneath the photoreceptor nuclear layer (mCherry) (Figure S1A). Müller glia processes, indicated by staining against glial glutamate/aspartate transporter (GLAST), extend over the photoreceptor nuclear layer, forming a notional outer limiting membrane (OLM). Inner and outer segment-like protrusions of photoreceptors are located right above this structure, visible in differential interference contrast (DIC) microscopy (Figure S1A). A significant number of mCherry^+^ cells co-stained with the cone-specific marker arrestin 3 (ARR3) and signs of cone maturation include expression of opsins of short (OPN1SW) and medium/long (OPN1M/LW) wavelength, though labelling was not restricted to outer segment-like protrusions but were found throughout the whole cell body (Figure S1B), indicative of an earlier state of differentiation.

To obtain an enriched fraction of transplantable photoreceptor cells, D200 HROs were dissociated into a single-cell suspension and underwent fluorescence-activated cell sorting (FACS) for mCherry^+^ cells, corresponding to ca. 65% of the alive (i.e. DAPI^-^) cell population (Figure S1C). Photoreceptor-identity of sorted mCherry^+^ cells was validated by immunocytochemistry against the pan-photoreceptor marker hRCVRN. Nevertheless, their characteristic photoreceptor architecture was lost, presenting a round morphology, likely attributed to shear stress during the steps of dissociation and FACS (Figure S1D).

### Characterization of degeneration status and Müller glia reactivity in hosts

Before delivery of donor cells into hosts, the mouse models were characterized to define extent of photoreceptor loss and potential Müller glia reactivity. In this study, two mouse models presenting different degeneration states were used to assess how the severity of retinal degeneration of the host might have an influence on the survival, incorporation and maturation of donor photoreceptors. Cpfl1 mice carry indel mutations in the phosphodiesterase 6C gene. They are characterized by dysfunctional cones and their subsequent loss, as indicated with staining against ARR3 (Figure 1A), but have remaining rod photoreceptors (Rhodopsin, RHO) and thus an outer nuclear layer (ONL) (Figure 1B), representing a case of mild retinal degeneration. In contrast, tg(Cpfl1/Rho^-/-^) mice, a strain resulting from crossing Cpfl1 to Rhodopsin knockout mice, present severe retinal degeneration with an almost complete loss of cones and rods and therefore also the ONL by the age of 16 weeks (Figure 1A, B).

**Figure 1.**
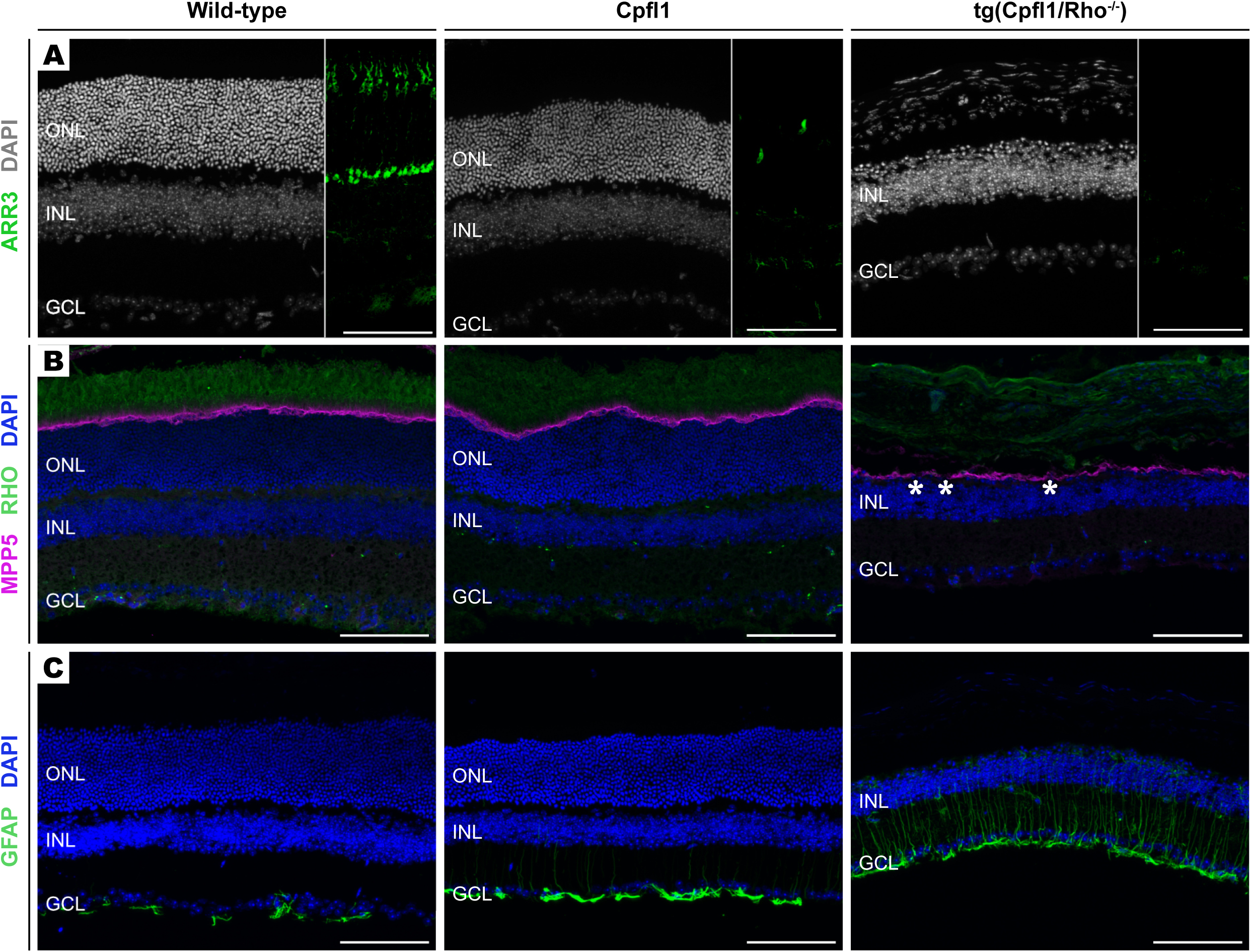
Cpfl1 and tg(Cpfl1/Rho^-/-^) mice present different states of photoreceptor degeneration and Müller Glia reactivity. (A) Cones, marked by Arrestin 3 (ARR3, green), are almost absent in Cpfl1 retinae, while the overall structure of the outer nuclear layer (ONL) is preserved, compared to its massive loss in the tg(Cpfl1/Rho^-/-^) mouse model. (B) Rods (rhodopsin^+^ (RHO), green) and outer limiting membrane (membrane protein palmitoylated 5^+^ (MPP5), magenta) are intact in wild-type and Cpfl1 mice, while tg(Cpfl1/Rho^-/-^) retinae show complete lack of rods and the outer limiting membrane appears compromised (asterisks). (C) Müller glia are reactive in both mouse models of retinal degeneration, as observed by the upregulation of glial fibrillary acidic protein (GFAP, green), with stronger expression in severely affected tg(Cpfl1/Rho^-/-^) mouse, contrary to wild-type retina, in which GFAP signal is restricted to astrocytes residing in the ganglion cell layer (GCL). INL: Inner nuclear layer. Scale bar: 100 μm.

Moreover, Müller glia, macroglia cells that span almost the entire retina, become reactive in degenerative states identified by the upregulation of glial fibrillary acidic protein (GFAP), a phenomenon often described as gliosis. GFAP is a cytoskeleton component of the cells which in homeostatic conditions (i.e. wild-type) is expressed solely by astrocytes residing in the ganglion cell layer (Figure 1C). In the tg(Cpfl1/Rho^-/-^) retina, GFAP is expressed throughout the Müller glia cell body, extending towards the apical processes, which is strongly pronounced when compared to the Cpfl1 retina, where GFAP expression is restricted to the level of the INL. Despite the lack of cones in Cpfl1 mice, its OLM appears to be continuous, while for the tg(Cpfl1/Rho^-/-^) mice the OLM appears to be sporadically disrupted or dimmer (asterisks), as indicated with staining against the membrane protein palmitoylated 5 (MPP5/PALS1), a scaffold component of the Crumbs complex (Figure 1B).

### Survival and clustering of transplanted cells are influenced by the host’s retinal degeneration severity

A suspension of 150,000 human donor photoreceptors was administered into the subretinal space of recipient Cpfl1 and tg(Cpfl1/Rho^-/-^) mice, followed by monthly intravitreal injections of the immunosuppressive triamcinolone acetonide starting with the transplantation. Donor cells, verified by staining against mCherry expression, survive up to 26 weeks post transplantation (wpt) (latest time point analyzed) in both mildly and severely degenerated retinal microenvironments (Figure 2A). However, graft volume differs between the two degeneration states. At 26 wpt, mean graft volume corresponds to 6,349,581 μm^3^ for Cpfl1 hosts, while in tg(Cpfl1/Rho^-/-^) to 2,340,832 μm^3^ (Figure 2B). Moreover, it appears that the degree of retinal degeneration severity influences the morphology of the graft. The images selected in figure 2A reflect retinal cross-sections with similar graft volume found for both models. In Cpfl1 hosts, donor photoreceptors tend to form multi-cellular clusters and are confined to a distinct retinal area, contrary to tg(Cpfl1/Rho^-/-^) hosts where they appear mostly singularized and are scattered, covering a larger part of the retina, but rarely packed in cell clusters (Figure 2C).

**Figure 2.**
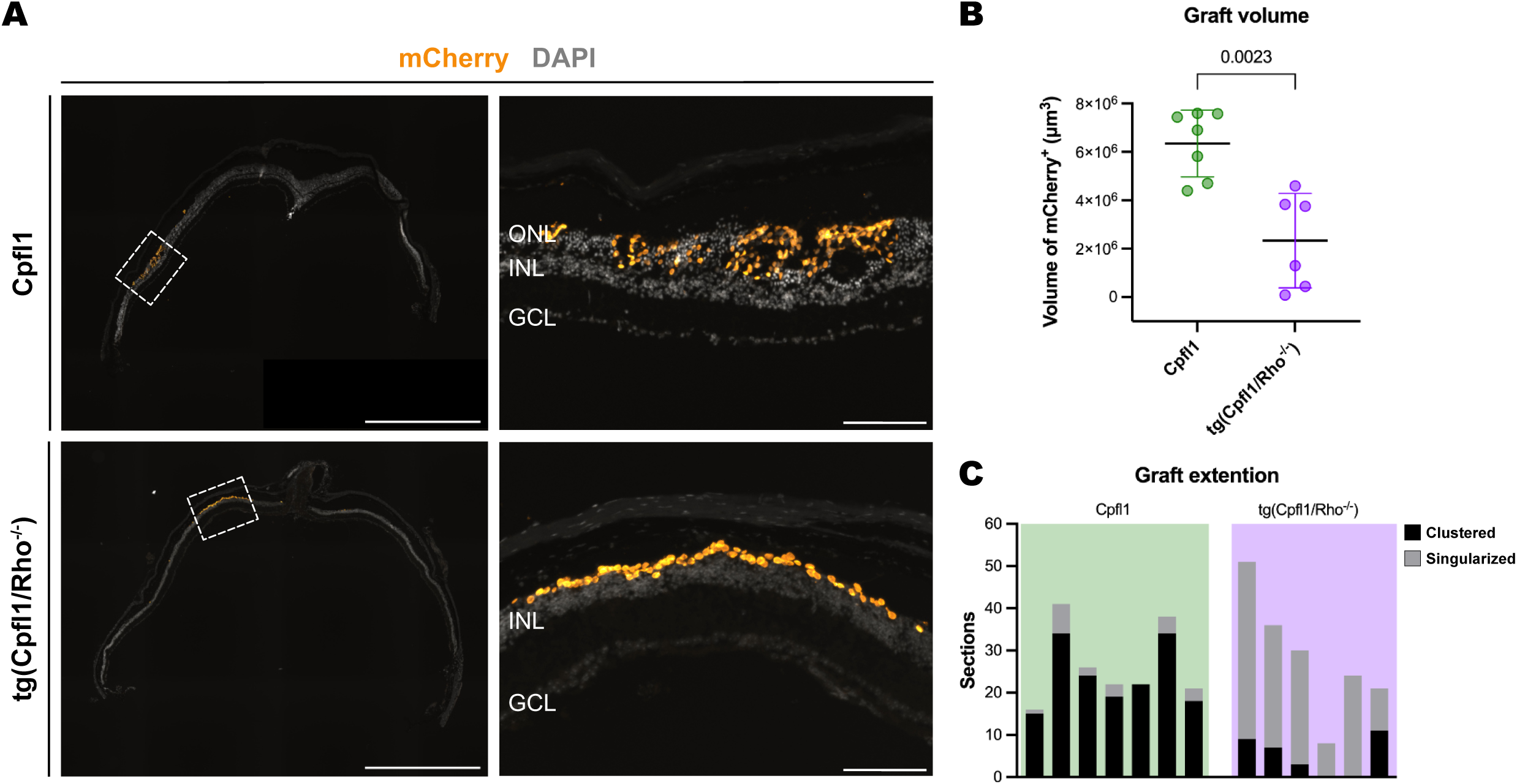
Grafts in Cpfl1 hosts show improved long-term survival, cover less area and appear more clustered in comparison to tg(Cpfl1/Rho^-/-^) hosts. (A) Grafts (mCherry, orange) in Cpfl1 (upper row) are more clustered and spread less compared to tg(Cpfl1/Rho^-/-^) hosts (lower row). Scale bars: 1000 μm (Insets: 100μm). (B) Graft volume is significantly bigger in Cpfl1 (6,349,581 ± 1,377,204 μm^3^) compared to tg(Cpfl1/Rho^-/-^) (2,340,832 ± 1,953,060 μm^3^) hosts. Each point represents one transplanted eye. Welch’s test, line at mean (SD). (C) Grafts mainly appear in multicellular clusters in Cpfl1 hosts, compared to tg(Cpfl1/Rho^-/-^) hosts, where donor photoreceptors are mostly found singularized. Each column represents one transplanted eye. ONL: outer nuclear layer; INL: inner nuclear layer; GCL: ganglion cell layer.

### Host Müller glia and transplanted photoreceptors interactions initiate at 3 wpt

Initial interactions between transplants and host cells were assessed at 3 wpt. At this time point, transplants still remain in the subretinal space in both models, with donor cells already more widely distributed in tg(Cpfl1/Rho^-/-^) in comparison to Cpfl1 hosts. In Cpfl1 mice, cellular ‘bridges’ between graft and host are formed at distinct locations as the host ONL forms contact points with the graft, and Müller glia processes grow extensively into the transplant as shown by staining against glutamine synthetase (GLUL) (Figure 3A, A’, upper row) and GFAP (Figure 3B, B’, upper row). At these contact points the OLM appears disrupted (Figure 3A, A’’ (asterisks)). In contrast, in tg(Cpfl1/Rho^-/-^) hosts, though the graft is laying in close proximity to the remaining host INL and is apposed to GLUL^+^ (Figure 3A, A’, bottom row) and GFAP^+^ (Figure 3B, B’, bottom row) processes, these processes seem not to grow into the graft and the OLM appears continuous (Figure 3A, A’’).

**Figure 3.**
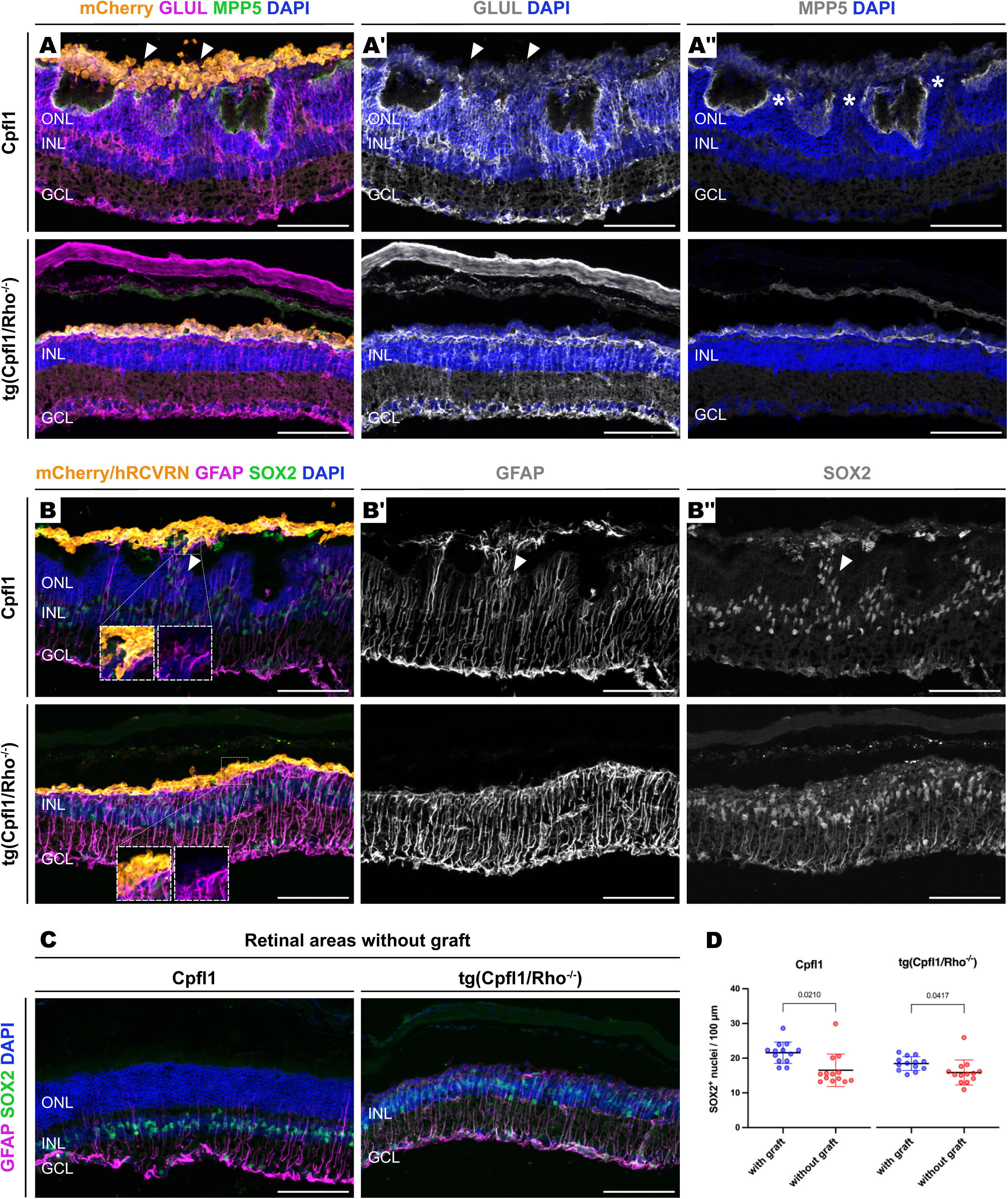
Interactions between host Müller glia and transplants at 3 wpt. (A) In Cpfl1 hosts Müller glia extent their processes (visualized by GLUL, A: magenta; A’: white) into the subretinally located graft (mCherry, orange) via distinct cellular ‘bridges’ (arrowheads), and the outer limiting membrane (marked with MMP5, A: green; A’’: white) shows disruptions at these contact points (A’’: asterisks). In tg(Cpfl1/Rho^-/-^) hosts (lower row), transplanted cells also remain in the subretinal space, but show no obvious interaction with host Müller glia processes, while the OLM appears undisrupted. (B) Staining for GFAP, a marker for reactive gliosis in Müller glia, confirms extensive growth of Müller glia processes into grafts in Cpfl1 but not tg(Cpfl1/Rho^-/-^) hosts (B: magenta, B’: white). Host Müller glia translocate SOX2^+^ nuclei passing the ONL towards interactive zones with transplants in Cpfl1 hosts (upper row, B: green; B’’: white), which is not observed in transplanted tg(Cpfl1/Rho^-/-^) hosts. (C) The reactive profile marked by GFAP (magenta) appears increased in graft-host interactive zones (B, B’) in comparison to retinal areas without transplant (C). In retinal areas without graft, SOX2^+^ nuclei reside in the INL and do not show signs of displaced nuclei. (D) Quantification of SOX2^+^ nuclei show a significant increase in areas with graft in comparison to areas without graft. n=3 per group, with 4-5 technical replicates per sample analyzed. Wilcoxon test, line at mean (SD). ONL: outer nuclear layer; INL: inner nuclear layer; GCL: ganglion cell layer; GLUL; glutamine synthetase; MPP5: membrane protein palmitoylated 5; RCVRN: recoverin; GFAP: glial fibrillary acidic protein; SOX2: SRY-box transcription factor 2. Scale bar: 100 μm.

Moreover, in Cpfl1 hosts, SOX2^+^ nuclei extend from the INL along these ‘bridges’ towards the graft (Figure 3B, B’’), compared to contralateral areas of the same eye where there is no graft present and SOX2^+^ expression is restricted to the INL and ganglion cell layer, where Müller glia and astrocytes typically reside, respectively (Figure 3C). In addition, quantitative analysis shows an increased number of SOX2^+^ nuclei in areas with graft compared to contralateral areas of the same retina (Figure 3D). Higher reactivity of Müller glia cells upon transplantation is also indicated by GFAP expression as the signal appears upregulated in the grafted areas compared to contralateral areas of the same eye (Figure 3B, B’, C). Overall, these results highlight a potential role for host Müller glia in mediating the initiation of graft incorporation.

On the other hand, when assessing other types of host cells, for example bipolar cells (Rod bipolar cells: Protein Kinase C alpha (PKCa) and cone bipolar cells: secretagogin (SCGN)), amacrine cells (Calretinin, CALB2), and horizontal cells (Neurofilament medium chain (NEFM)) (Figure S2), no arborizations or processes are observed interacting with donor photoreceptors at 3 wpt.

### Increased structural integration of transplants at 10 and 26 wpt

By 10 wpt the grafts in both Cpfl1 and tg(Cpfl1/Rho^-/-^) hosts show increased incorporation within the remaining neural retina (Figure 4A). Large parts of transplants in Cpfl1 hosts were no longer located in the subretinal space, but instead structurally integrated into the host ONL (Figure 4A, upper row, arrowheads), Host Müller glia processes - observed with staining against GLUL - enwrap photoreceptors within the transplants (Figure 4A, A’, upper row, arrowheads), while the OLM – marked by MPP5 - appears disrupted in regions of graft integration (Figure 4A’’, upper row, asterisks). Signs of structural integration were also observed in tg(Cpfl1/Rho^-/-^) hosts, where single donor cells showed interactions with host Müller glia processes (Figure 4A, A’, lower row, arrowheads), while also the OLM was no longer continuous, but showed several disruptions (Figure 4A, A’’, lower row, asterisks).

**Figure 4.**
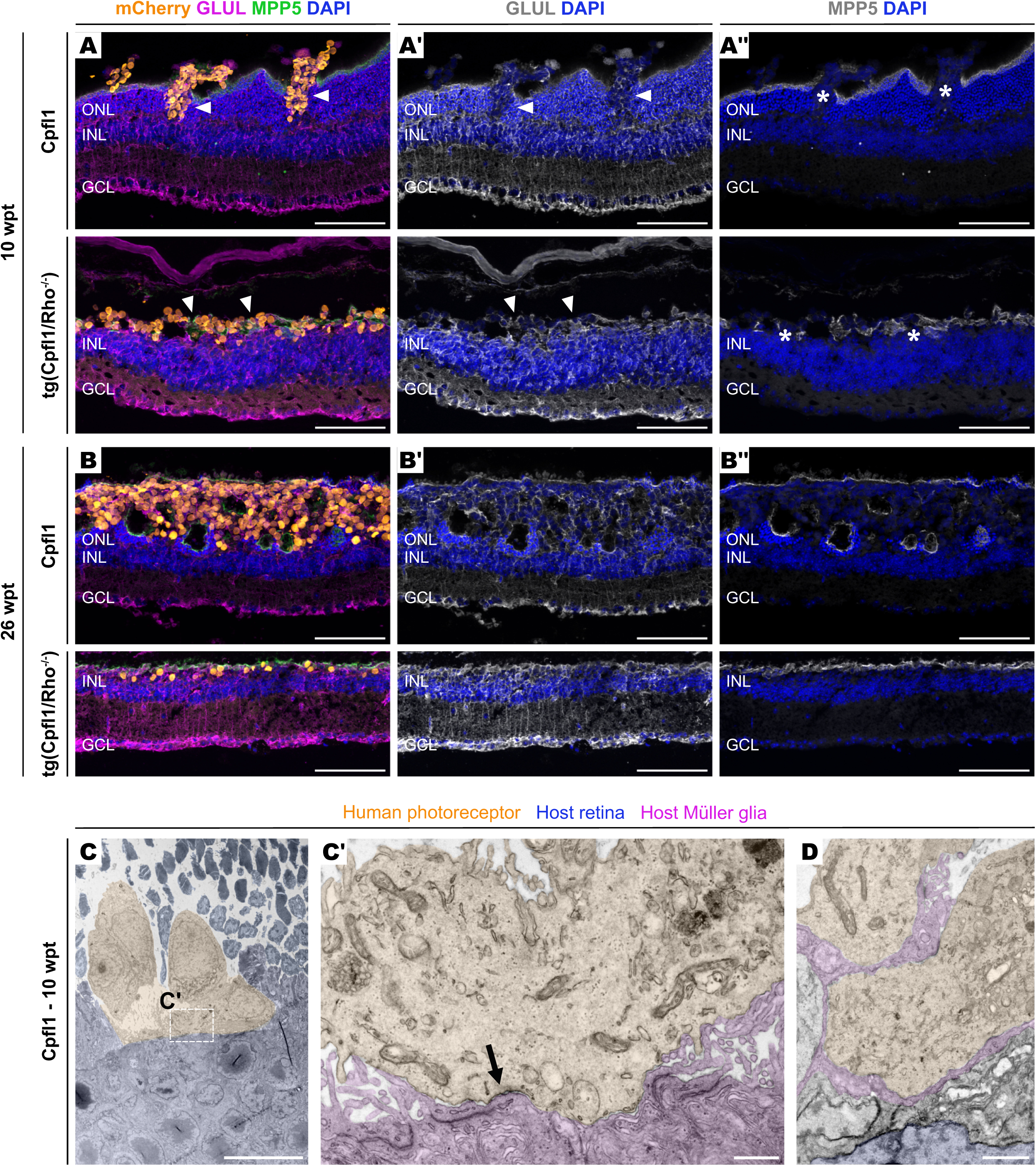
Increased integration of human photoreceptors at 10 and 26 wpt. (A) At 10 wpt, human transplants (mCherry, orange) show partial incorporation into the ONL of Cpfl1 hosts (upper row), with enwrapment of donor photoreceptors by Müller glia processes marked with GLUL (A: magenta, A’: white). At the sites of partial incorporation, the OLM (MPP5, A: green, A’’: white) is disrupted (marked by asterisks). First signs of enwrapment of donor photoreceptors by Müller glia processes are also observed at 10 wpt in tg(Cpfl1/Rho^-/-^) hosts (A, A’, lower row, some marked by arrowheads), while the OLM shows a disrupted appearance (A’’, lower row). At 26 wpt, many donor photoreceptors appear deeper incorporated into the retina of tg(Cpfl1/Rho^-/-^) hosts with further increased interaction with Müller glial processes (B, B’), while the OLM is mainly closed and continuous, with donor cell bodies located basally (B’’, lower row). Transmission electron microscopy images show close proximity of donor photoreceptors (pseudo-colored orange) and host Müller glia (pseudo-colored magenta) in Cpfl1 hosts at 10 wpt (C, C’, D) including formation of tight junctions (marked by an arrow). ONL: outer nuclear layer; INL: inner nuclear layer; GCL: ganglion cell layer; GLUL: glutamine synthetase; MPP5: membrane protein palmitoylated 5.Scale bars: 100 μm (A, A’, A’’, B, B’, B’’), 10 μm (C), 500 nm (C’), 1 μm (D).

Structural integration of donor photoreceptors into the host retina was confirmed for Cpfl1 hosts by ultrastructural electron microscopy analysis, where tight junctions were formed between human photoreceptors and host Müller glia (Figure 4C, C’ (arrow)), and Müller glia processes were identified in close contact to nascent inner segments of donor photoreceptors (Figure 4D).

By 26 wpt, transplants appear to be fully incorporated within the neural retina of Cpfl1 hosts (Figure 4B, upper row), with Müller glia processes enveloping cells within the graft (Figure 4B’, upper row), while the OLM appears re-sealed and continuous with cell bodies of donor photoreceptors located basally of it (Figure 4B’’, upper row). Interestingly, also in tg(Cpfl1/Rho^-/-^) hosts, where there are no endogenous photoreceptors left at this time point, donor cells appear increasingly incorporated, with several donor cells interacting with Müller glia processes (Figure 4B, B’, lower row). Also, the OLM appears continuous, with donor cell bodies located basally (Figure 4B’’, lower row).

### Maturation of donor cells is affected by the degenerative microenvironment

By 10 wpt, when host Müller glia largely interact with donor photoreceptors, first signs of maturation are also distinct. Immunohistochemical staining against human-specific mitochondria (hMITO) reveals that donor photoreceptors in Cpfl1 hosts have formed single cellular substructures attached to their cell bodies, resembling inner segments typically found in maturing photoreceptors (Figure 5A, left panel). These bulbi colocalize with expression of recoverin (RCVRN), a pan-photoreceptor marker, and are often apically polarized into the subretinal space towards the RPE. Additionally, RCVRN^+^ processes are formed by donor photoreceptors towards the host INL, reminiscent of axonal processes (Figure 5A, left panel, marked by arrowheads). In contrast, donor photoreceptors in tg(Cpfl1/Rho^-/-^) hosts rarely developed inner segment- or axon-like structures at 10 wpt, with mitochondria mainly distributed throughout the cell body (Figure 5A, right panel). At 26 wpt, donor photoreceptors in Cpfl1 hosts show signs of further maturation with high numbers of apical hMITO^+^ inner segment-like protrusions and, additionally, outer segment-like structures, as indicated by staining against opsin 1 long/medium wave sensitive (OPN1L/MW), attached to hMITO^+^ protrusions (Figure 5B, left panel). On the contrary, maturation of donor photoreceptors in severely degenerated hosts did not resemble the case of Cpfl1 hosts, as hMITO and OPN1L/MW^+^ extensions are rarely observed (Figure 5B, right panel). Quantification of the number of hMITO puncti in Cpfl1 and tg(Cpfl1/Rho^-/-^) hosts at 26 wpt reflects that a significantly larger number of transplanted photoreceptors formed inner segment-like structures, particularly with apical polarization, within Cpfl1 retinae compared to the severely degenerated tg(Cpfl1/Rho^-/-^) retina (Figure 5C).

**Figure 5.**
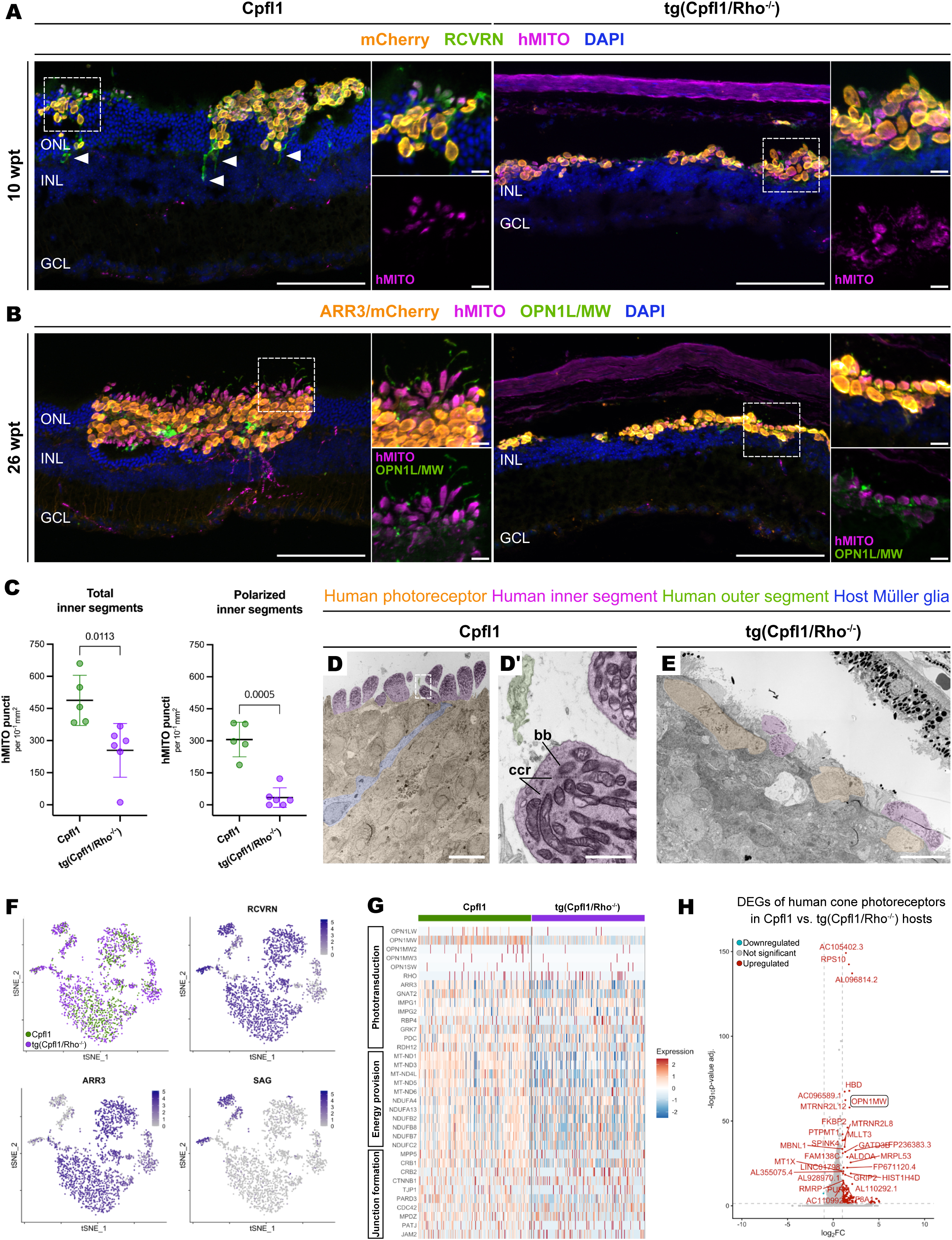
Maturation of donor photoreceptors is pronounced in Cpfl1 compared to tg(Cpfl1/Rho^-/-^) hosts. (A, left panel) Staining against recoverin (RCVRN, green) and human-specific mitochondrial content (hMITO, magenta) reflects nascent formation and polarization of inner segments of donor photoreceptors (mCherry, orange), as well as formation of axonal processes towards the INL (marked by arrowheads) in Cpfl1 hosts at 10 wpt. (A, right panel) Donor photoreceptors in tg(Cpfl1/Rho^-/-^) hosts rarely show hMITO^+^ inner-segment-like protrusions or axonal processes at 10 wpt. (B, left panel) At 26 wpt, donor photoreceptors (ARR3/mCherry, orange) proceed to mature further in Cpfl1 hosts generating many apical oriented inner-segment-like protrusions (hMITO, magenta) and additionally forming outer segment-like extensions identified by staining against cone-specific opsins (OPN1L/MW, green). (B, right panel) In tg(Cpfl1/Rho^-/-^) few hMITO^+^ inner segment-like protrusions are attached to donor photoreceptors at 26 wpt, while OPN1L/MW^+^ puncti are rare and apically oriented. Scale bars: 100 μm (Insets: 10 μm). (C) Quantification of hMITO^+^ puncti per 10^-1^ mm^2^ as a proxy for inner segment formation. Transplanted photoreceptors in Cpfl1 hosts form significantly more total (487.8 ± 117.1 vs. 254.2 ± 125.3 puncti) as well as polarized inner segments in comparison to tg(Cpfl1/Rho^-/-^) hosts (306.2 ± 80.6 vs. 34.3 ± 45.9 puncti). Each point represents one transplanted eye. Values normalized to graft area. Welch’s test, line at mean (SD). (D, D’) Transmission electron microscopy images showing a cluster of human photoreceptors (pseudo-colored orange) integrated into the retina of Cpfl1 hosts at 26 wpt with apical mitochondria-enriched inner segments (pseudo-colored magenta), nascent outer segments (pseudo-colored green) and basal body (bb) with connecting cilium rootlets (ccr). (E) Some cell bodies of donor photoreceptors (pseudo-colored orange) in tg(Cpfl1/Rho^-/-^) hosts show integration or partial integration into the retinal tissue, with nascent, apically oriented inner segments, that contain few mitochondria. Scale bars: 10 μm (Inset: 1 μm). (F) t-SNE plots from scRNA-seq analysis indicates that the donor cells are photoreceptors given the high expression of the pan-photoreceptor gene RCVRN, with the majority of donor cells expressing cone-specific ARR3 and some expressing rod-specific SAG. (G) Heatmap reflecting expression of phototransduction-, energy provision- and junction formation-related genes in 100 representative donor cells isolated from Cpfl1 and tg(Cpfl1/Rho^-/-^) hosts. Overall, expression of these genes associated with maturation is higher in Cpfl1 hosts. (H) Volcano plot of differentially expressed genes (DEGs) in donor cells from Cpfl1 and tg(Cpfl1/Rho^-/-^) hosts show upregulation of several genes associated with mature photoreceptors in Cpfl1 hosts. ONL: outer nuclear layer; INL: inner nuclear layer; GCL: ganglion cell layer; ARR3: Arrestin 3; OPN1L/MW: opsin 1 long/medium wave sensitive; SAG: S-antigen visual arrestin.

Ultrastructural analysis of experimental retinae using electron microscopy at 26 wpt confirms extensive incorporation of transplanted human photoreceptors as clusters into Cpfl1 retinae. These are accompanied by mature inner segments which are enriched in mitochondria with a basal body (bb) neighbouring to connecting cilium rootlets (ccr) as well as nascent outer segments (Figure 5D, D’). On the contrary, transplanted human photoreceptors in tg(Cpfl1/Rho^-/-^) retinae appear singularized and only partially incorporated into the host tissue, albeit those are also accompanied by nascent inner segments (Figure 5E). However, inner segments appear less structured with less mitochondria content and no outer segment-like structures could be detected. Generally, nuclei and inner segments of human photoreceptors were significantly larger than of mouse photoreceptors, and human nuclei did not form the characteristic inverted architecture with centrally condensed heterochromatin as seen in mouse rod photoreceptors (Figure S3A). Additionally, ultrastructural analysis of grafts 26 wpt revealed the formation of mature ribbon synapses flanked by synaptic vesicles in Cpfl1 hosts (Figure S3A, A’). On the contrary, only rarely naïve synaptic-like structures were detected within donor photoreceptors in tg(Cpfl1/Rho^-/-^) retinae (Figure S3B, B’).

### Expression of maturation markers by donor photoreceptors is increased in Cpfl1 vs. tg(Cpfl1/Rho^-/-^) hosts

To evaluate the molecular signature of human photoreceptors after transplantation into the distinct degenerative phenotypes, biopsies containing donor cells were isolated from transplanted retinae and analyzed via single-cell RNA-sequencing (scRNA-seq) at 10 wpt (Figure S4A). Evaluation of key markers confirmed that donor cells represent photoreceptors (Figure 5F, Figure S4B; exemplary photoreceptor-markers: RCVRN, CRX), with the majority (ca. 80%) corresponding to cones (Figure 5F; exemplary cone marker: ARR3) and the most remaining cells expressing rod genes (Figure 5F; exemplary rod marker: SAG). Extremely rarely, Müller glia cells or bipolar cells showed up in the donor cell population, as examined by the expression of the markers vimentin (VIM) and carbonic anhydrase 10 (CA10), respectively (Figure S4C, D). This confirms that Müller glia or bipolar cell signals observed in close contact or within the transplants derived from the host, and were not introduced by the transplanted cell suspension. Moreover, expression of genes involved in the processes of phototransduction, energy provision, and junction formation – all characteristics of a mature photoreceptor – is higher in transplants within Cpfl1 mice compared to tg(Cpfl1/Rho^-/-^) hosts (Figure 5G; heatmap represents gene expression of 200 donor photoreceptors down-sampled per condition), underlining an overall improved maturation status of donor photoreceptors in the milder degenerative microenvironment. Indeed, when assessing differentially expressed genes (DEGs) of the cone fraction from the two hosts, OPN1MW appears to be one of the highest significantly upregulated genes of donor photoreceptors within Cpfl1 recipients (Figure 5H).

### Clustered donor cells do not show improved incorporation and maturation in tg(Cpfl1/Rho^-/-^) hosts

Donor photoreceptors showed a wider distribution and singularization after transplantation into tg(Cpfl1/Rho^-/-^) mice, while in corresponding Cpfl1 hosts donor cells formed larger cell clusters with signs of improved integration and maturation. To evaluate whether the number of clustered grafts can be increased and thus interactions with host Müller glia as well as integration and photoreceptor maturation can be improved, a suspension of 500,000 donor cells (instead of 150,000 cells) was administered in both models and assessed 10 wpt. Even though that donor cells had an increased propensity of forming clusters in the tg(Cpfl1/Rho^-/-^) recipients, they appeared mostly isolated from the rest of the host retina (Figure 6A, B, right panels). Compared to the nicely structured and polarized inner segments in Cpfl1 hosts, indicated by hMITO and hRCVRN expression, such bulbi were still rarely observed in the severe model of retinal degeneration (Figure 6).

**Figure 6.**
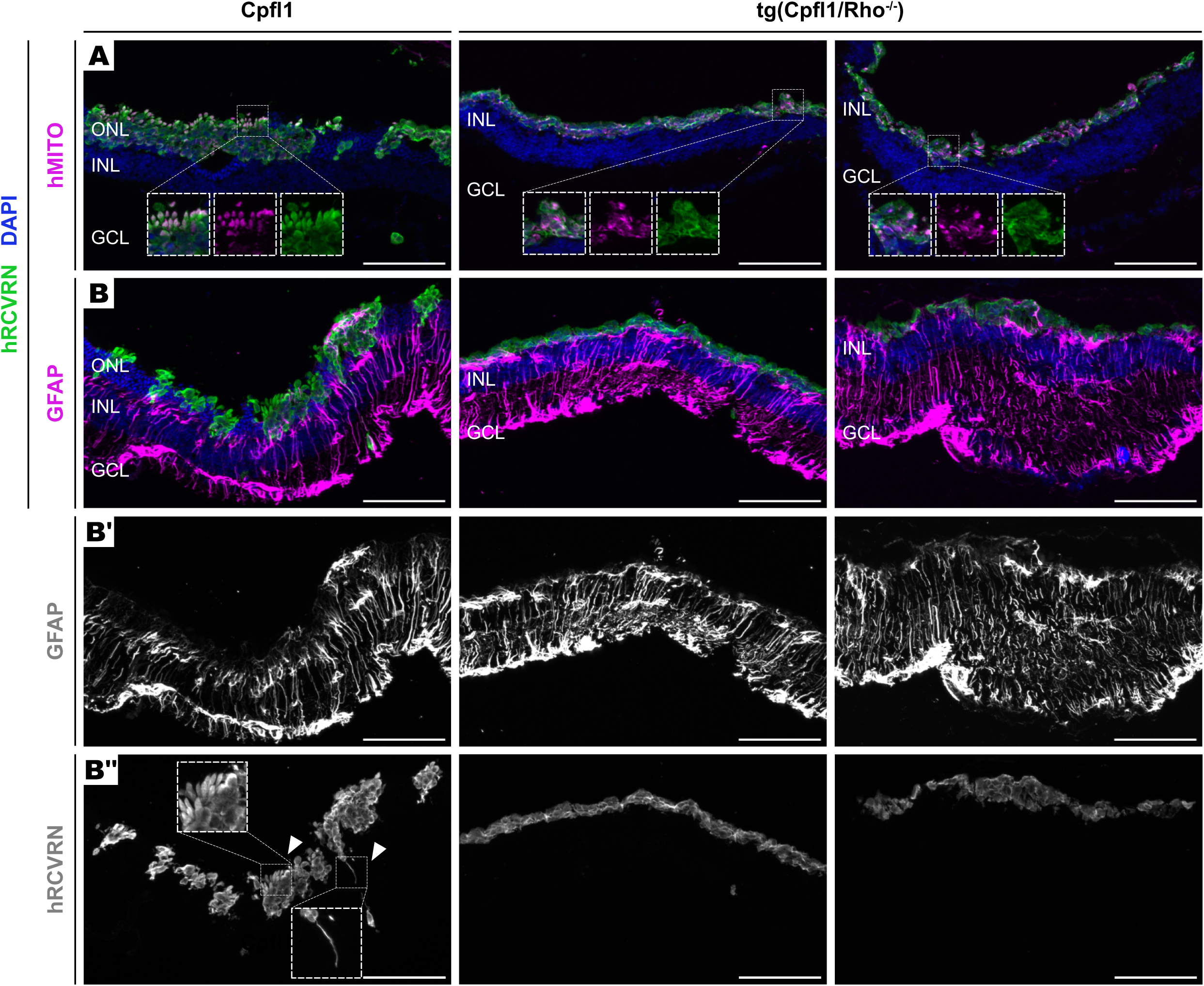
Clustering of transplanted photoreceptors does not suffice for their maturation. (A) Following transplantation of 500,000 cells, donor photoreceptors (hRCVRN, green) are clustered in both Cpfl1 and tg(Cpfl1/Rho^-/-^) recipients at 10 wpt, but hMITO^+^ (magenta) inner segments are formed and apically polarized much more in the Cpfl1 retina. (B) Host Müller glia (marked with GFAP, B: magenta; in B’: white) interact vastly with the graft in Cpfl1 contrary to tg(Cpfl1/Rho^-/-^) hosts. (B’’) Donor photoreceptors expressing hRCVRN (white) generate apical inner segments and basal axonal processes in Cpfl1 but not tg(Cpfl1/Rho^-/-^) hosts. ONL: outer nuclear layer; INL: inner nuclear layer; GCL: ganglion cell layer; hRCVRN: human-specific recoverin; hMITO: human-specific mitochondrial content; GFAP: glial fibrillary acidic protein. Scale bar: 100 μm.

Moreover, interactions between host Müller glia and donor photoreceptors remain sparse in the tg(Cpfl1/Rho^-/-^) mouse (Figure 6B, B’, right panel). On the contrary, donor photoreceptors have partially incorporated into the host ONL of Cpfl1 hosts and are widely enwrapped by host Müller glia processes (Figure 6B, B’, left panel). Staining against human-specific recoverin (hRCVRN) highlights the mature structure of donor photoreceptors in Cpfl1 hosts, as both, segments and neurites, can be found, whereas such structures are mainly absent from tg(Cpfl1/Rho^-/-^) hosts (Figure 6B, B’’). These findings further support that donor-host interactions are crucial for their subsequent incorporation and maturation, and that the morphology of the graft – whether donor photoreceptors are in clusters or singularized – does not influence these processes in tg(Cpfl1/Rho^-/-^) hosts.

### Transplantation at an earlier stage of progressive degeneration results in improved incorporation and maturation of donor photoreceptors as well as remodelling of host bipolar cells

Hypothesizing that the retina of Cpfl1 hosts might provide a better scaffold due to the remaining ONL, 150,000 donor cells were transplanted into 4 weeks old tg(Cpfl1/Rho^-/-^) mice - henceforth referred to as tg(Cpfl1/Rho^-/-^)^early^ - a time point when approximately 90% endogenous photoreceptors, and thus an ONL, are still present (Figure S5). Qualitative immunohistological assessment of grafts 26 wpt in tg(Cpfl1/Rho^-/-^)^early^ hosts revealed a clustering phenotype consisted of mature photoreceptors similar to those observed in Cpfl1 hosts (Figure 7A). Inner segments, defined by staining against hMITO and hRCVRN, appear analogous in shape to those formed in Cpfl1 hosts and are largely polarized towards the subretinal space/RPE (Figure 7B). Host Müller glia GLUL^+^ processes enveloped donor photoreceptors and a nascent OLM was formed (MPP5) (Figure 7C). Interestingly, endogenous photoreceptors, of which 90% were still present at the time of transplantation, were completely lost in tg(Cpfl1/Rho^-/-^)^early^ host at 26 wpt despite integration and maturation of donor photoreceptor clusters.

**Figure 7.**
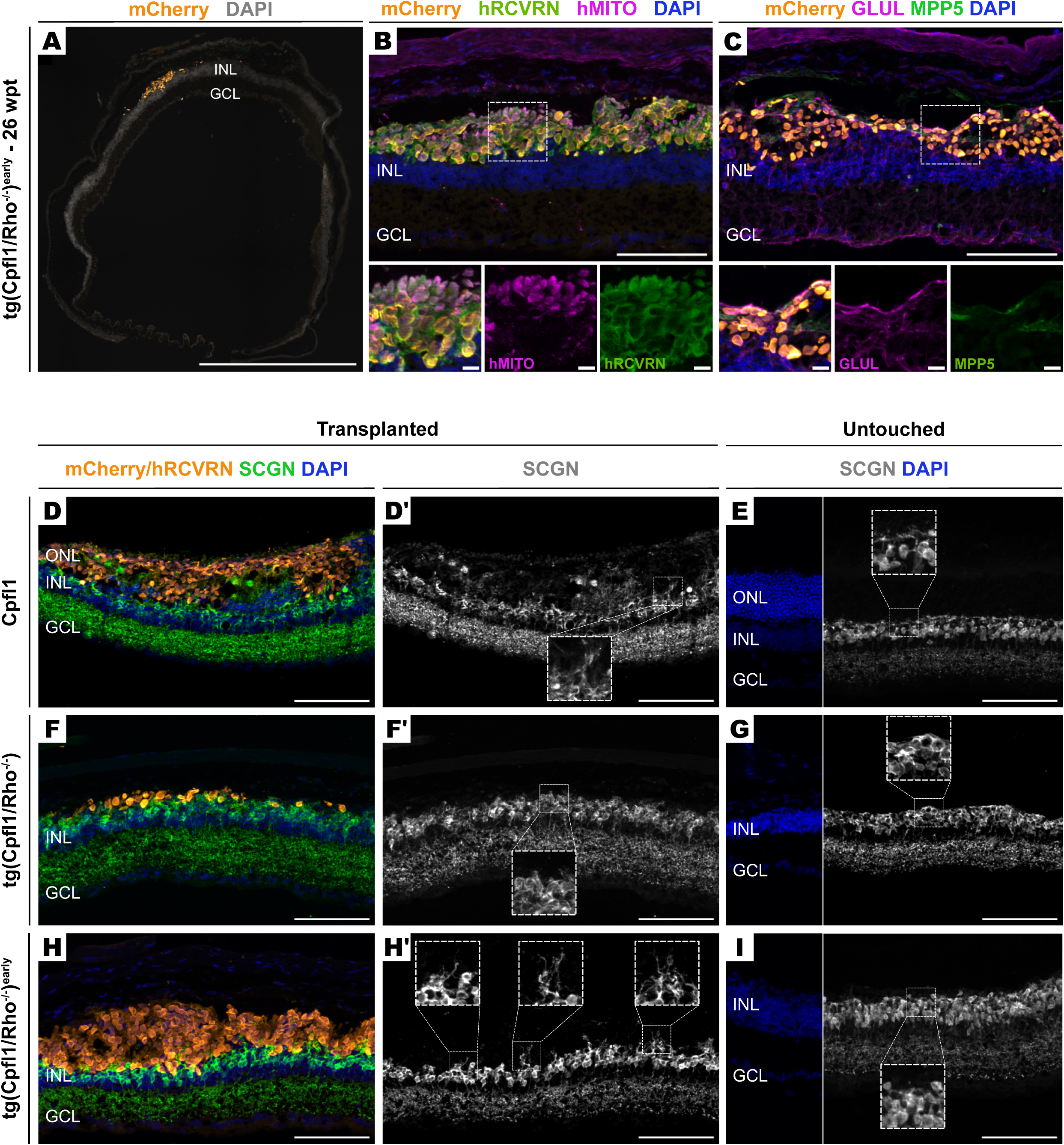
Photoreceptor transplants in tg(Cpfl1/Rho^-/-^)^early^ hosts show improved clustering and maturation as well as incite dendrite regrowth of host bipolar cells. (A) Donor photoreceptors (mCherry, orange; hRCVRN, green) appear clustered in tg(Cpfl1/Rho^-/-^)^early^ hosts and (B) generate polarized inner segments (hMITO, magenta). (C) Host Müller glia (GLUL, magenta) envelop the donor cells in tg(Cpfl1/Rho^-/-^)^early^ hosts and a nascent outer limiting membrane (MPP5, green) is formed. (D) In Cpfl1 hosts, cone bipolar cells grow dendrite processes (SCGN, green; in D’: white) towards/into the transplant (mCherry/hRCVRN, orange). (E) Apical dendrite processes, though shorter, are also present in the untouched Cpfl1 retina. (F, F’, G) tg(Cpfl1/Rho^-/-^) retinae do not form dendrite outgrowths, neither with or without photoreceptors transplanted at a late degeneration stage. While host cone bipolar cells in untouched retinae retract their dendrites in tg(Cpfl1/Rho^-/-^)^early^ at 26 wpt, SCGN^+^ dendrite outgrowths are visible in areas containing transplants. ONL: outer nuclear layer; INL: inner nuclear layer; GCL: ganglion cell layer; hRCVRN; human-specific recoverin; hMITO: human- specific mitochondrial content; GLUL: glutamine synthetase; MPP5: membrane protein palmitoylated 5; SCGN: Secretagogin. Scale bar: 100 μm (Insets: 10 μm).

In Cpfl1 hosts, cone bipolar cells (marked by secretagogin – SCGN) grow dendritic processes deeply into structurally integrated transplants (Figure 7D, D’), with extended length in comparison to cone bipolar cells in non-transplanted retinae (Figure 7E). In age-matched untouched tg(Cpfl1/Rho^-/-^) and tg(Cpfl1/Rho^-/-^)^early^ retinae (i.e. 42 and 30 weeks old, respectively), host bipolar cells have fully retracted their dendrites most likely due to complete degeneration of the host ONL (Figure 7G, I). In tg(Cpfl1/Rho^-/-^) retinae transplanted at late-stage, growth of cone bipolar dendritic processes remained low in regions with or without donor cells (Figure 7F, F’). In contrast, in tg(Cpfl1/Rho^-/-^)^early^ hosts, bipolar cells undergo remodelling and their dendrites re-grow towards the graft (Figure 7H, H’), similar as in Cpfl1 hosts.

## DISCUSSION

Pre-clinical studies from several laboratories have provided first evidence for successful transplantation of human photoreceptors isolated from PSC-derived retinal organoids including some functional improvements in mouse models of photoreceptor loss.^13^ Retinal degenerative diseases are characterized by extremely high genetic (up to now more than 500 genes with disease-causing mutations identified),^14^ and phenotypic heterogeneity, with diverse complex pathologies described, ranging from early to late onset and slow to fast photoreceptor death. While photoreceptor replacement represents a potential generalized disease-agnostic treatment approach, insights into whether and to what extent such highly diverting disease environments affect photoreceptor transplants have not been systematically assessed. In this study we utilized two mouse models showing distinct levels of photoreceptor degeneration to investigate in which ways the underlying retinal degenerative status influences transplanted human photoreceptors in regard to survival, structural integration, and maturation by employing histological, ultra-structural and transcriptome analysis. We observed that photoreceptor transplants in the severely degenerated retina survive, cluster, and mature to a much lesser extent compared to the milder model of retinal degeneration. Interestingly, transplantation into the severe mouse model at an early disease stage allows structural integration and maturation of transplants despite continuous loss of endogenous photoreceptors.

Human donor photoreceptors, enriched from iPSC-derived retinal organoids at D200, were subretinally transplanted into mild (Cpfl1) or severe tg(Cpfl1/Rho^-/-^) degenerative hosts. While in both disease environments donor cells showed long-term survival for up to 26 wpt, the latest time point examined, grafts showed significant differences in their distribution within the subretinal space: in Cpfl1 recipients donor photoreceptors mainly form multi-cellular clusters, whereas in tg(Cpfl1/Rho^-/-^) mice they cover a larger part of the retina but appear singularized. Quantification of combined nuclei volume of donor cells within the two mouse models revealed significantly bigger volumes in Cpfl1 hosts arguing for improved survival. Singularized distribution was also observed by Collin and colleagues when transplanting human ESC-derived CRX^+^ cells into the severely degenerated Pde6brd1 mouse line.^1^ On the contrary, in a study by Ribeiro and colleagues,^4^ in which 500,000 human cone photoreceptors were transplanted into the severely degenerated Rd1 mouse, donor cells appear clustered. Similar findings were observed by Procyk and colleagues,^7^ in which the Aipl1^-/-^ mouse model - presenting complete ONL degeneration – was used. Indeed, larger grafts were also observed in our study after transplantation of 500,000 cells into the late-stage mouse model, however, this did not improve incorporation and maturation of donor photoreceptors.

When comparing graft morphologies between models at the three different time points examined - 3, 10, and 26 wpt - it is apparent that donor cells overtime cluster more in Cpfl1 hosts. Rarely, rosette-like structures were formed by donor photoreceptors or by the underlying host photoreceptors at 26 wpt. This feature of photoreceptor re-organization resembles tubulations observed in retinal degeneration speculated to form upon sustained photoreceptor or RPE damage,^16^ however, the reason behind their formation in the context of photoreceptor replacement therapy is currently unknown. Donor cells in tg(Cpfl1/Rho^-/-^) hosts remain rather dispersed and stretched throughout the length of the retina. We hypothesize that the interphotoreceptor matrix, the extracellular matrix coating the subretinal space, might also undergo remodelling proportional to the extent of photoreceptor degeneration, which could in turn influence the clustering phenotype. This is further supported by our results regarding transplantations in the early degeneration stage of tg(Cpfl1/Rho^-/-^) hosts, where only 10% of endogenous photoreceptors are lost at the time of transplantation and in which the donor photoreceptors remain clustered at 26 wpt. Host cells might also participate to these morphological differences as at 3 wpt they contact the graft in Cpfl1 hosts, and the graft from that point onwards becomes more compacted and spatially restricted. Notably, the delivery of more donor cells in transplantations, 500,000 vs. 150,000, does potentiate more interactions between them, a parameter which is tunable.

Interestingly, scRNA-seq analysis revealed that more cones than rods are present in the graft population at 10 wpt, a selectivity which remains to be investigated, given that similar proportion of cones and rods are observed in retinal organoids produced by our standardized protocol.^11^ Additionally, donor cones predominantly differentiated towards the medium wavelength cone subtype. Adjusting the number and proportion of cell types of administered cells are elements for consideration for tailoring potential cell therapies to the needs of patients (e.g., patients with achromatopsia would require a cone-rich transplantation, while retinitis pigmentosa patients would require rod-rich transplants and at later stages with complete blindness would require both photoreceptor types).

At 3 wpt host Müller glia expand their processes towards the graft, co-aligning with our previous observations when transplanting human iPSC-derived cones into Cpfl1 hosts,^2^ and translocate their nucleus from the INL to ONL. An increase of the SOX2^+^ nuclei is also found in the transplanted areas, compared to non-transplanted areas of the same retina, an event which is not model-specific, occurring both for Cpfl1 and tg(Cpfl1/Rho^-/-^) hosts. Such phenomena suggest gliogenesis, also observed by Sudharsan and colleagues,^17^ the process of glial proliferation which is characterized with interkinetic nuclear migration. As these responses where restricted in the retinal areas containing grafts, they indicate a Müller glia-specific response in mediating graft incorporation. However, it remains unclear whether this translocation is true interkinetic nuclear migration or a whole cell body migration event. Importantly, even though Müller glia pre-transplantation appear to have a reactive state in both Cpfl1 and tg(Cpfl1/Rho^-/-^) retinae, given the observed GFAP upregulation, this does not appear to be inhibitory towards host-graft interactions, as often suggested due to the formation of a gliotic seal observed in retinal degeneration. ^18–20^ Furthermore, even though Cpfl1 retinae are characterized with a continuous OLM, it appears that Müller glia and the OLM per se do not hinder incorporation but rather are required for graft incorporation, as previously noted.^2^ Overall, these results suggest that host Müller glia are the first responders mediating interactions with the graft, a dynamic process entailing nuclei/cell body disposition and possibly cell proliferation. The grafts appear fully incorporated by 26 wpt in the Cpfl1 hosts, and largely incorporated into the tg(Cpfl1/Rho^-/-^) hosts. Taking into consideration that grafts at 3 wpt in the severely degenerated hosts remained isolated from the rest of the neural retina, these observations suggest delayed incorporation and as such differential incorporation kinetics into the host. Understanding how early host-graft interactions potentiate the long-term graft survival and functional integration efficiency might be key for treatment of patients with late-stage pathology.

Differences were also observed regarding the maturation of transplanted photoreceptors within the two degeneration models, at both the phenotypic as well as molecular level. We observed formation of polarized inner and outer segment-like structures in the transplanted photoreceptors in Cpfl1 hosts, a morphology closely reminiscent of a mature photoreceptor. On the contrary, less and disorganized segments are found in tg(Cpfl1/Rho^-/-^) hosts where both hMITO^+^ and OPN1L/MW^+^ signals were diffusely distributed within the transplants, underlining an influence of retinal degeneration severity on *in vivo* maturation of donor photoreceptors. The morphological differences were recapitulated by scRNA-seq analysis, showing increased expression of genes related to maturation of donor photoreceptors within Cpfl1 hosts including phototransduction, energy metabolism, and junction formation. Others have reported partial functionality rescue in severely degenerated retinae,^3,4,7,21,22^ despite donor photoreceptors providing also less signs of full maturation, such as the ones observed here within the mild model. It is therefore of interest to assess to what extent graft maturation influences functionality post-transplantation, a correlation analysis which was beyond the scope of this study.

Advanced maturation of donor cells within Cpfl1 host might result from the closer/stronger interaction with host Müller glia cells (see above). Indeed, in retinal development and physiology Müller glia partake in assembly, as well as proper function and integrity of photoreceptors. ^23,24^ Thus, deciphering the cellular and molecular factors facilitating interactions between host Müller glia and donor photoreceptors might identify targets for improving graft integration in future studies. Interestingly, increasing the number of donor photoreceptors to 500,000 cells, which allowed for better/more contacts between the graft and host tissue in the tg(Cpfl1/Rho^-/-^) hosts, was not sufficient for further photoreceptor maturation. However, while interactions with Müller glia cells appear to be a main factor for donor photoreceptor polarization and maturation, other physiological components might also play important roles in this process. This might entail the surrounding subretinal matrix, as transplanted photoreceptors at the tg(Cpfl1/Rho^-/-^)^early^ stage showed better morphology including inner segment formation. Other factors might include synapse formation between donor photoreceptors with second order neurons, or interactions with RPE cells. The latter is of importance, as certain pathologies are also characterized by RPE degeneration, which could in turn damage further transplanted photoreceptors, as maintenance and health of light-sensitive cells highly depends on the integrity of this epithelium. By investigating therefore different parameters of the diseased retinal microenvironment, we hope to improve the degree of incorporation and maturation of donor photoreceptors following transplantation into different states of retinal degeneration.

Thus, these findings provide evidence that the development and physiological status of photoreceptor transplants is significantly influenced by the recipient environment and should be taken into account in the development of photoreceptor replacement therapies.

## MATERIALS & METHODS

### Maintenance of human iPSCs and differentiation in retinal organoids

The iPSC AAVS1::CrxP H2BmCherry line ^2,10^ used for the generation of human retinal organoids (HROs) was kindly provided by Olivier Goureau (Institute de la Vision, Paris, France) and maintained at the Stem Cell Facility of the Center for Molecular and Cellular Bioengineering (CMCB, TU Dresden). iPSCs were cultured in mTeSR™ 1 medium (STEMCELL Technologies, Cologne, Germany, #85850) on vitronectin-coated plates (Gibco, Dreieich, Germany, #A400458) and passaged using ReLeSR^TM^ (STEMCELL Technologies, Cologne, Germany, #100-0484) at 37 °C. Differentiation of iPSCs into HROs was performed using a previously optimized protocol.^25^ Briefly, on D0 undifferentiated iPSCs at 60-80% confluency were washed twice with Phosphate-buffered saline (PBS) and incubated with ReLeSR^TM^ for 2 minutes (min) at 37 °C. Cells were resuspended in DMEM/F-12 + GlutaMAX^TM^ (Gibco, Dreieich, Germany, #31331028), collected using wide-bore pipette tips, and centrifuged for 3 min at 700 rpm at room temperature (RT). The cell pellet was placed in ice and resuspended in N2B27 medium [1:1 DMEM/F-12 + GlutaMAX^TM^: Neurobasal medium (Gibco, Dreieich, Germany, #21103-049), 1% B27 with Vitamin A (Gibco, Dreieich, Germany, #17504044), 0.5% N2 (Gibco, Dreieich, Germany, #17502048), 1% penicillin-streptomycin (Sigma-Aldrich, Taufkirchen, Germany, #P0781), 50 μM β-mercaptoethanol (Gibco, Dreieich, Germany, #31350010)]. Growth factor-reduced Matrigel (Corning, Berlin, Germany, #354230) was added to the cell solution and allowed to polymerize for ca. 7 min. The Matrigel was dispersed in N2B27 medium, and aggregates were seeded in low-attachment 6-well plates (Thermo Scientific, Dreieich, Germany, #174945). Fresh medium was added on D3, and on D5 neuroepithelial cysts were transferred to Matrigel-coated 6-well plates at a 1:2 ratio. Fresh media was added on D7 and D9, followed by a full media change on D11. On D13, adherent cysts were washed twice with DMEM/F-12 and detached using Dispase (STEMCELL Technologies, Cologne, Germany, #07923). After a 15 min incubation at 37 °C, the cysts were mechanically detached and collected into 15 ml tubes. After two washing steps with DMEM/F12, the cysts were transferred to ultra-low attachment Petri dishes (Thermo Scientific, Dreieich, Germany, #174945) containing B27 medium [DMEM/F12 GlutaMAX^TM^, 2% B27 without Vitamin A (Gibco, Dreieich, Germany, #12-587-010), 1% penicillin-streptomycin, 1% Non-Essential Amino Acids (Gibco, Dreieich, Germany, #11140035), 0.1% amphotericin B (Gibco, Dreieich, Germany, #15290026)]. A half medium change was done the day after, and then every 2-3 days. On D25, the medium was replaced with B27/FBS medium [B27 media with 10% Fetal Bovine Serum (Gibco, Dreieich, Germany, #A5256801)] plus 0.3 μM of EC23 (Tocris, Wiesbaden-Nordenstadt, Germany, #4011). Between D30 and D40, retinal epithelial evaginations were manually isolated using surgical tweezers. On D100, the HROs were cultured in RM2 medium (DMEM/F12 GlutaMAX^TM^, 1% N2, 10% FBS, 1% penicillin-streptomycin, 0.1% amphotericin B) plus 0.3 μM of EC23. After 20 days (D120) EC23 was completely removed and medium was changed every 2-3 days until D200.

### FACS of mCherry^+^ cells

D200 HROs were dissociated into a single-cell suspension following our established protocol.^12^ In brief, the HROs were incubated with 20 U/mL papain for 1.5 hours (h) and then mechanically triturated using a narrow fire-polished glass pipette and processed according to the manufacturer’s instructions (Papain Dissociation System, Worthington). The cell pellet was then resuspended in Magnetic-activated cell sorting (MACS) buffer (0.5% BSA, 2 mM EDTA in PBS) which was filtered through a 35 μm mesh reaching a final concentration of ca. 7 Mio cells/mL. Samples were stored on ice and then sorted with either AriaFusion, AriaII or AriaIII, using a 100 μm nozzle at 4 °C. Before isolation of mCherry^+^ cells based on the fluorescent intensity, debris and doublets were removed by adjusting the forward-scatter and side-scatter areas. 4′,6-diamidino-2-phenylindole (DAPI) (Germany, AppliChem GmbH, Darmstadt, Germany, #A1001.0010) staining was applied to remove dead cells.

### Animals

Adult Cpfl1-mutant (12-19 weeks of age) and tg(Cpfl1/Rho^-/-^)-mutant (15-18 weeks of age) mice were used as recipients for cell-suspension transplantation. The colonies were maintained in the CRTD animal facility on a 12 h light/12 h dark cycle with *ad libitum* access to food and water. After intervention, gel food was provided to the animals and regular health check-ups were performed.

### Optical coherence tomography

The set up for OCT was used as described previously. ^26^ Briefly, mice were anesthetized with an intraperitoneal injection of 100 mg/mL ketamin (Serumwerk Bernburg AG, Bernburg, Germany, #13690.00.00) and 1 mg/mL medetomidine (Orion Pharma, Hamburg, Germany, #32457.00.00) mixture, and a drop of 2.5% phenylephrine/0.5% tropicamide solution (University Clinic Pharmacy, Dresden, Germany) was added to the eye for 2-3 minutes to dilate the pupil. The excess was removed and hydrating gel (Bausch + Lomb, Berlin, Germany, #40612.00.00), was added before placing a glass coverslip on the cornea. The OCT scanner was placed in parallel to the eye, and cross-sections of the central retina were acquired every 4 μm. After the final scan anaesthesia was immediately reversed by intraperitoneal injection of 5 mg/mL Antisedan (Orion Pharma, Hamburg, Germany, #23554.00.00). Recordings were processed in ImageJ (NIH).

### Transplantations

Post FACS, mCherry^+^ cells were resuspended in MACS buffer (150,000 cells/μL or 500,000 cells/μL) and 1 μL was delivered into the subretinal space of anesthetized recipient mice with a Hamilton syringe mounted to a micromanipulator as described in Tessmer et al. (2023).^12^ Following transplantation and on a monthly basis, 1 μL of Triamcinolone acetonide suspension (80 μg/μL in NaCl, University Clinic Pharmacy, Dresden, Germany) was delivered intravitreally. Anaesthesia was immediately reversed.

### Immunohistochemistry and immunocytochemistry

For immunohistochemistry, transplanted mice were euthanized and eyes were enucleated and fixed in 4% paraformaldehyde in PBS for 1 h at 4 °C. The muscle, cornea, lens and optic nerve were dissected away, and the eyes were cryopreserved in a 30% w/v sucrose solution overnight at 4 °C. The tissue was then embedded in optimal cutting medium NEG50 (Epredia, Dreieich, Germany, #6502), and 12 μm cross-sections were collected on Superfrost^TM^Plus Adhesion Microscope Slides (Epredia, Dreieich, Germany, #J1800AMNZ), and dried in a heating platform. Slides were first hydrated with PBS for 30 min and then incubated with blocking solution (0.3% Triton X-100, 1% BSA, 5% donkey solution in PBS) for 1h at RT, before incubation with primary antibodies overnight at 4 °C. The next day, 3 x 10 min washes with PBS performed, followed by a 90 min incubation with respective secondary antibodies, and another 3 x 10 min washes with PBS. Lastly, the slides were dried and mounted in Aqua-Poly/Mount (Polysciences, Hirschberg an der Bergstrasse, Germany, #18606-20) and let fully dry. Primary and secondary antibodies used are listed in Tables 1 and 2, respectively, in supplemental information. All secondary antibodies were used in a 1:1000 dilution.

**Table 1.**
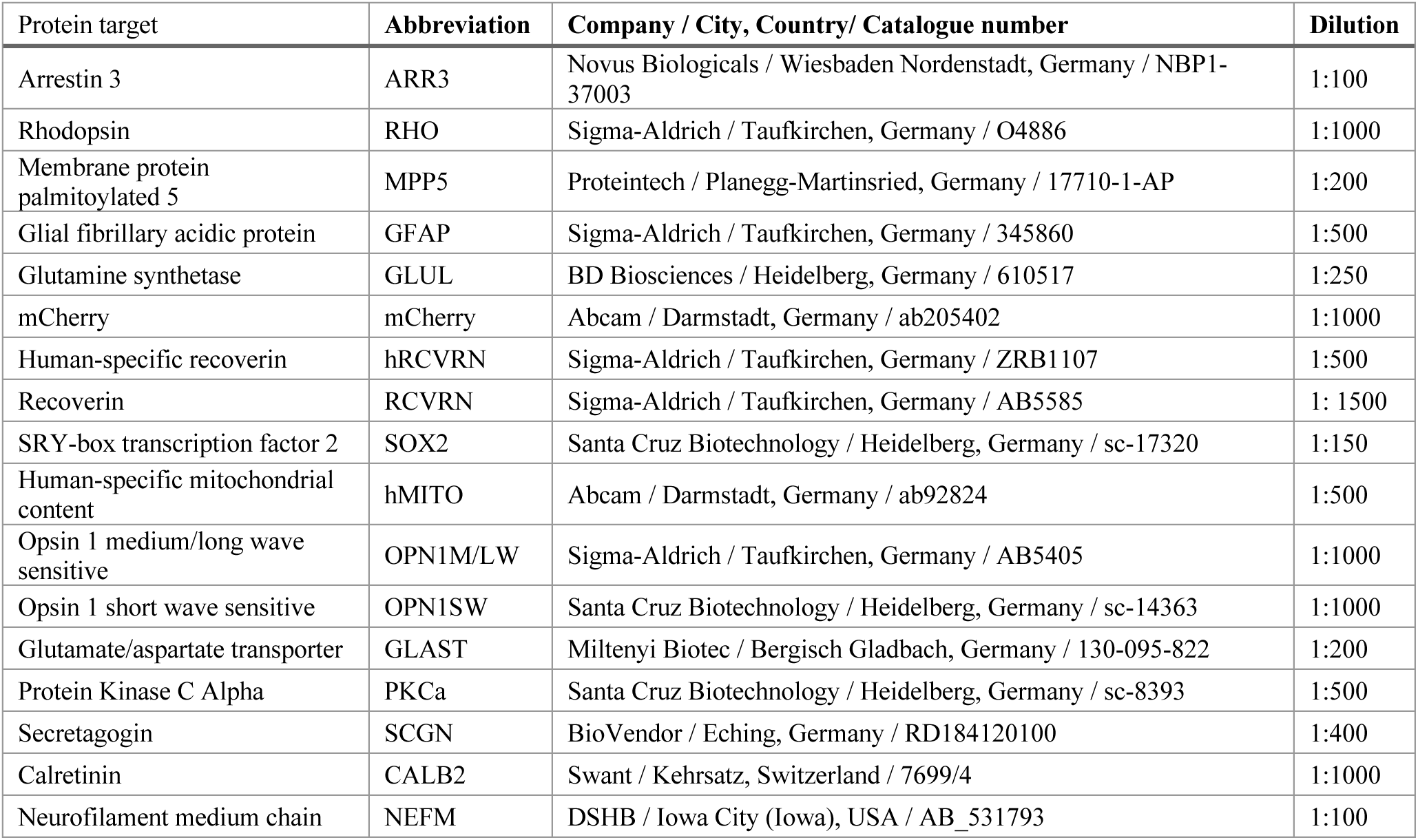
Primary antibodies.

**Table 2.** Secondary antibodies.

| Product | Company / City, Country / Catalogue number |
| --- | --- |
| Cy<sup>TM</sup>5 AffiniPure<sup>®</sup> Donkey Anti-Rabbit IgG (H+L) | Jackson ImmunoResearch Laboratories / Hamburg, Germany / 711-175-152 |
| Cy<sup>TM</sup>3 AffiniPure<sup>®</sup> Donkey Anti-Rabbit IgG (H+L) | Jackson ImmunoResearch Laboratories / Hamburg, Germany / 711-165-152 |
| Alexa Fluor<sup>®</sup> 488 AffiniPure<sup>®</sup> Donkey Anti-Rabbit IgG (H+L) | Jackson ImmunoResearch Laboratories / Hamburg, Germany / 711-545-152 |
| Cy<sup>TM</sup>5 AffiniPure<sup>®</sup> Donkey Anti-Mouse IgG (H+L) | Jackson ImmunoResearch Laboratories / Hamburg, Germany / 715-175-150 |
| Cy<sup>TM</sup>3 AffiniPure<sup>®</sup> Goat Anti-Mouse IgG (H+L) | Jackson ImmunoResearch Laboratories / Hamburg, Germany / 115-165-146 |
| Cy<sup>TM</sup>2 AffiniPure<sup>TM</sup> Donkey Anti-Mouse IgG (H+L) | Jackson ImmunoResearch Laboratories / Hamburg, Germany / 715-225-150 |
| Cy<sup>TM</sup>3 AffiniPure<sup>®</sup> Donkey Anti-Chicken IgY (IgG) (H+L) | Jackson ImmunoResearch Laboratories / Hamburg, Germany / 703-165-155 |
| Cy<sup>TM</sup>5 AffiniPure<sup>®</sup> Donkey Anti-Goat IgG (H+L) | Jackson ImmunoResearch Laboratories / Hamburg, Germany / 705-175-147 |
| Cy<sup>TM</sup>3 AffiniPure<sup>®</sup> Donkey Anti-Goat IgG (H+L) | Jackson ImmunoResearch Laboratories / Hamburg, Germany / 705-165-147 |
| Alexa Fluor<sup>®</sup> 488 AffiniPure<sup>®</sup> Donkey Anti-Goat IgG (H+L) | Jackson ImmunoResearch Laboratories / Hamburg, Germany / 705-545-147 |
| Alexa Fluor<sup>®</sup> 488 AffiniPure<sup>®</sup> Donkey Anti-Sheep IgG (H+L) | Jackson ImmunoResearch Laboratories / Hamburg, Germany / 713-225-147 |
| Alexa Fluor<sup>®</sup> 647 AffiniPure<sup>®</sup> Donkey Anti-Rat IgG (H+L) | Jackson ImmunoResearch Laboratories / Hamburg, Germany / 712-605-153 |
| DAPI | Sigma-Aldrich / Taufkirchen, Germany / D9542 |

### Transmission electron microscopy (TEM)

Mice were sacrificed and transplanted eyes were dissected to remove the muscle, cornea, lens and optic nerve and the rest was fixed with 4 % formaldehyde in 100 mM phosphate buffer for 2 h at RT followed by a transfer to 1 % FA/PBS for storage. The eyes were then further dissected under a fluorescence stereomicroscope (Leica MZ10F) to isolate labelled regions of interest (ROI) in blocks of less than 1 mm size. These blocks were further processed for TEM according to a modified protocol developed for serial block face SEM which involves osmium tetroxide (OsO4), thiocarbohydrazide (TCH), and again OsO4 to generate enhanced membrane contrast. ^2,25,27^ In brief, samples were postfixed overnight in modified Karnovsky fixative (2 % glutaraldehyde/2 % formaldehyde in 100 mM PBS, pH 7.4), followed by washes in PBS and water and post-fixation in a 2 % aqueous OsO4 solution containing 1.5 % potassium ferrocyanide and 2 mM CaCl2 (30 min on ice), washes in water, 1 % TCH in water (20 min at RT), washes in water and a second osmium contrasting step in 2% OsO4/water (30 min on ice).

Samples were washed and en-bloc contrasted with 1% uranyl acetate/water for 2 h on ice, washed in water, and dehydrated in a graded series of ethanol/water mixtures (30%, 50%, 70%, 90%, 96%, 20 min each), followed by three changes in pure water-free ethanol on molecular sieve (30 min each). Samples were infiltrated into the epon substitute EMBed 812 (resin/ethanol mixtures: 1:3, 1:1, 3:1 for 1 h each, followed by pure resin overnight and for 5 h) and finally cured at 65 °C overnight. To identify ROIs in the blocks semithin sections (1 μm) were prepared with a Leica UC6 ultramicrotome (Leica Microsystems, Wetzlar, Germany) using glass knifes, and stained with toluidine blue/borax for histological analysis. Once located, the ROIs were cut ultrathin (70 nm) using a diamond knife (Diatome, Nidau, Switzerland), the sections were collected on formvar-coated slot grids, and stained with 2 % aqueous uranyl acetate and with lead citrate.^28^ Mounted sections were analyzed with a JEM 1400Plus transmission electron microscope (JEOL, Freising, Germany) at 80 kV and images and montages were taken with a Ruby digital camera (JEOL, Freising, Germany).

### Image acquisition and processing

Samples were imaged using a Zeiss Apotome Imager Z2 (Germany). For evaluation of graft size, full Axioscan images were acquired with a Zeiss Axio Scan.Z1 (Germany). Images were processed using ImageJ2 (version 2.3.0/1.53s) and Affinity Designer (version 1.10.4). Images shown were generated using the Max Intensity Projection algorithm on z-stacks of 1 μm intervals in Image J2. Graft volume was quantified based on the fluorescent CRX-mCherry signal post immunohistochemistry from every fourth serial retinal cross-section throughout each transplanted eye. Affinity Designer was used for the generation of graphical abstract and schematic representation.

### Isolation and sample preparation of transplanted cells for single-cell transcriptomics

Retinae were dissected away from freshly enucleated eyes. The transplanted areas were isolated with a puncher tool under RFP channel of fluorescent microscope to detect the donor cells. Samples from 3-4 experimental eyes from each group were pooled together into one sequencing experiment to increase yield. The areas were dissociated first enzymatically with papain in a 11 mM D-Glucose in PBS solution, followed by mechanic trituration using a wide fire-polished glass pipette. The cells were carefully resuspended and visually inspected under a light microscope to determine cell concentration and quality. The single-cell suspensions were carefully mixed with the reverse transcription mix and were then loaded on the 10X Genomics Chromium system (Prague, Czech Republic) in a Chromium Single Cell G Chip (#PN-1000120) along with gel beads from Chromium Next GEM Single Cell 3’ v3.1 Gel Bead Kit v3.1 (#PN-1000122). Following the guidelines of the 10x Genomics Chromium Single Cell Kit v3.1 user manual (User guide ID: CG000315) and using the Chromium Next GEM Single Cell 3’ GEM Kit v3.1 (#PN-1000123), the droplets were directly subjected to reverse transcription, the emulsion was broken and cDNA was purified using Dynabeads MyOne Silane (#PN-2000048). After the amplification of cDNA with 11/12 cycles, 0.6x purification was perform with SPRIselect beads (Beckman Coulter, Krefeld, Germany, #B23319) to enrich cDNA fragments (>400 bp) and a quality control was performed on the Fragment Analyzer (using the DNF-473 NGS Fragment Kit/DND474 NGS High Sensitivity Kit, Agilent, Waldbronn, Germany). The 10X Genomics single cell RNA-seq library preparation - involving fragmentation, dA-Tailing, adapter ligation and a 11/13 cycles indexing PCR – was performed based on the manufacturer’s protocol using 10x Genomics Library Construction Kit (#PN-1000190). After quantification, the libraries were sequenced on an Illumina Novaseq6000 in paired-end mode (R1/R2: 100 cycles; I1/I2: 10 cycles), generating ∼140-360 million fragment pairs.

### Τranscriptome analysis

The raw sequencing data was processed with the count command of the Cell Ranger software (v6.1.2) provided by 10x Genomics (https://www.10xgenomics.com/). Since samples were a mix of human and mouse cells, Cell Ranger was run using a combined reference (GRCh38 and mm10, 2020-A, https://www.10xgenomics.com) to separate the 10x cell barcodes by species. Cell Ranger indices were built following the instructions provided by 10x Genomics (https://www.10xgenomics.com/support/software/cellranger/downloads/cr-ref-build-steps, 2020-A). Analysis of the 10x Genomics single-cell RNA-sequencing data was performed on Seurat (version 5.3.0)^29^ in Rstudio (version 2025.09.0+387). Quality control was performed to filter for cells of poor quality by adjusting the parameters for percentage of mitochondrial genes expressed and number of transcripts present. Data was then normalized, scaled, and integrated. Expression of selected genes was done using the DoHeatmap package (version 5.4.0) from Seurat. The top 30 DEGs, with an adjusted p-value < 0.05, between the cone fractions of Cpfl1 and tg(Cpfl1/Rho^-/-^) were visualized in the volcano plot using the package ggplot2 (version 4.0.3).

### Statistics

Statistical tests were performed in GraphPad Prism (version 10.6.1). For the graft volume analysis, as well as the hMITO^+^ puncti quantifications, the values passed the Kolmogorov-Smirnov normality test and then analyzed with the Unpaired t test with Welch’s correction. The values derived from the SOX2^+^ nuclei quantifications were not normally distributed and therefore the non-parametric Wilcoxon test was used. Values portrayed with mean (SD).

## DATA AVAILABILITY STATEMENT

We are in the process of safely storing the sequencing data on the GHGA platform (German Human Genome-Phenome Archive; https://www.ghga.de) upon manuscript publication.

## ACKNOWLEDGMENTS

The authors would like to thank the staff of the Light Microscope, Flow Cytometry, Dresden-concept Genome Center, Stem Cell Engineering and Animal facilities of the CRTD/CMCB, for their assistance with Axioscan imaging, cell sorting, scRNA-seq, cell culturing and mouse maintenance, respectively. Further thanks go to Sabine Klaussner, Jochen Hentschel, and Susanne Fergusson for technical assistance. This work was supported by grants from the Deutsche Forschungsgesellschaft (DFG): AD 375/7-1 and AD 375/7-2 (within the SPP2127; to M.A.), AD 375/9-1 (to M.A.), Foundation Fighting Blindness: TA-RM-0522-0824-TUD-TRAP to M.A.). Co-funded by the ERASMUS+ programme of the European Union: 2024-1-EL01-KA131-HED-000210895 (to A.M.). Views and opinions expressed are however those of the authors only and do not necessarily reflect those of the European Union or the State Scholarships Foundation (IKY). Neither the European Union nor the granting authority can be held responsible for them.

## AUTHOR CONTRIBUTIONS

M.P. designed and performed experiments, analyzed and visualized data, and wrote the manuscript, K.T. participated in experiment design and data curation, T.K. performed electron microscopy imaging and contributed in the methodology part of the manuscript, J.H. performed optical coherence tomography, A.M. and C.P. participated in data analysis, B.C.S.M. performed cell culture experiments and contributed in the methodology part of the manuscript, F.R. supervised the analysis of the RNA-seq data, M.A. conceptualized the project, supervised, reviewed and edited the manuscript.

---

**Figure S1. D200 human retinal organoids generate transplantable photoreceptors.** (A) D200 HROs present a laminated structure with Müller glia nuclei (SOX2, magenta) residing underneath photoreceptor nuclei (mCherry, orange), while their processes (GLAST, green) extend apically forming a notional OLM. (B) Photoreceptors present their distinctive morphology as seen with the cytosolic expression of cone arrestin (ARR3, orange), and express short wave-length sensitive opsins (OPN1SW, magenta) as well as medium/long-wavelength opsins (OPN1L/MW, green). (C) FACS allows the isolation and enrichment of mCherry^+^ cells (orange) - gating indicates the target cell fraction. (D) Post FACS mCherry^+^ cells lose their distinctive photoreceptor morphology and round up with the majority expressing the pan-photoreceptor marker hRCVRN (green), and some expressing the cone-specific marker ARR3 (magenta). OLM: outer limiting membrane; ONL: outer nuclear layer; INL: inner nuclear layer; SOX2: SRY-box transcription factor 2; GLAST: glutamate/Aspartate Transporter 1; DIC: Differential Interference Contrast; ARR3: arrestin 3; OPN1L/MW: opsin 1 long/medium wave sensitive; OPN1SW: opsin 1 short wave sensitive; FACS: fluorescence-activated cell sorting; FSC: forward scatter; SSC: side scatter; hRCVRN: human-specific recoverin. Scale bars: 100 μm (Insets: 25 μm).

**Figure S2. Bipolar, amacrine and horizontal cells do not interact with donor photoreceptors at 3 wpt.** (A) Labelling against Protein Kinase C alpha (PKCa, magenta) and Secretagogin (SCGN, green) shows no interactions between bipolar cells and donor photoreceptors (mCherry, orange; hRCVRN, green) at 3 wpt. (B) Labelling against Calretinin (CALB2, magenta) and Neurofilament medium chain (NEFM, green) shows no interactions between amacrine/horizontal cells, respectively, and donor photoreceptors (mCherry, orange) at 3 wpt. ONL: outer nuclear layer; INL: inner nuclear layer; GCL: ganglion cell layer; hRCVRN: human-specific recoverin (hRCVRN). Scale bar: 100 μm.

**Figure S3. TEM reveals ribbon synapse formation by donor photoreceptors in mildly degenerated host.** (A, A’) Example of a mature synapse containing a ribbon and surrounding vesicles generated by an integrated donor photoreceptor at 26 wpt. (B - identical to Figure 5E, B’) Very rarely premature synaptic structures are identified in tg(Cpfl1/Rho-/-) hosts at 26 wpt.

**Figure S4. Evaluation of donor cell population through scRNA-seq analysis.** (A) Pipeline of isolation of human transplanted cells for scRNA-seq. (B) t-SNE plot showing Cone-rod homeobox (CRX) transcript expression in donor cell population. (C) t-SNE plot showing rare Vimentin (VIM) transcript expression (specific for macroglia/Müller glia) in donor cell population. (D) t-SNE plot showing rare Carbonic anhydrase 10 (CA10) transcript expression (specific for bipolar cells) in donor cell population.

**Figure S5. tg(Cpfl1/Rho^-/-^) mice present age-dependent retinal degeneration.** Progressive decrease of the ONL (marked in red) in tg(Cpfl1/Rho^-/-^) mice at ages 4 and 16 weeks is visible by (A) *in vivo* optical coherence tomography (OCT) imaging, and (B) differential interference contrast (DIC) imaging and DAPI staining on cross-sections. Images were acquired from central retinal cross-sections areas adjacent to optic nerve. ONL: outer nuclear layer; INL: inner nuclear layer; GCL: ganglion cell layer. Scale bar: 50 μm.

## REFERENCES

1. Collin, J., Zerti, D., Queen, R., Santos-Ferreira, T., Bauer, R., Coxhead, J., Hussain, R., Steel, D., Mellough, C., Ader, M., et al. (2019). CRX Expression in Pluripotent Stem Cell-Derived Photoreceptors Marks a Transplantable Subpopulation of Early Cones. Stem Cells 37, 609–622. 10.1002/stem.2974.

2. Gasparini, S.J., Tessmer, K., Reh, M., Wieneke, S., Carido, M., Völkner, M., Borsch, O., Swiersy, A., Zuzic, M., Goureau, O., et al. (2022). Transplanted human cones incorporate into the retina and function in a murine cone degeneration model. 10.1172/JCI154619.

3. Mandai, M., Fujii, M., Hashiguchi, T., Sunagawa, G.A., Ito, S., Sun, J., Kaneko, J., Sho, J., Yamada, C., and Takahashi, M. (2017). iPSC-Derived Retina Transplants Improve Vision in rd1 End-Stage Retinal-Degeneration Mice. Stem Cell Reports 8, 69–83. 10.1016/j.stemcr.2016.12.008.

4. Ribeiro, J., Procyk, C.A., West, E.L., O’Hara-Wright, M., Martins, M.F., Khorasani, M.M., Hare, A., Basche, M., Fernando, M., Goh, D., et al. (2021). Restoration of visual function in advanced disease after transplantation of purified human pluripotent stem cell-derived cone photoreceptors. Cell Reports 35, 109022. 10.1016/j.celrep.2021.109022.

5. Thomas, B.B., Rajendran Nair, D.S., Rahimian, M., Hassan, A.K., Tran, T.-L., and Seiler, M.J. (2025). Animal models for the evaluation of retinal stem cell therapies. Progress in Retinal and Eye Research 106, 101356. 10.1016/j.preteyeres.2025.101356.

6. Matsuyama, A., Kalargyrou, A.A., Smith, A.J., Ali, R.R., and Pearson, R.A. (2022). A comprehensive atlas of Aggrecan, Versican, Neurocan and Phosphacan expression across time in wildtype retina and in retinal degeneration. Sci Rep 12, 7282. 10.1038/s41598-022-11204-w.

7. Procyk, C.A., Melati, A., Ribeiro, J., Liu, J., Branch, M.J., Delicata, J.D., Tariq, M., Kalarygrou, A.A., Kapadia, J., Khorsani, M.M., et al. (2025). Human cone photoreceptor transplantation stimulates remodeling and restores function in AIPL1 model of end-stage Leber congenital amaurosis. Stem Cell Reports, 102470. 10.1016/j.stemcr.2025.102470.

8. Chang, B., Hawes, N.L., Hurd, R.E., Davisson, M.T., Nusinowitz, S., and Heckenlively, J.R. (2002). Retinal degeneration mutants in the mouse. Vision Research 42, 517–525. 10.1016/S0042-6989(01)00146-8.

9. Santos-Ferreira, T., Völkner, M., Borsch, O., Haas, J., Cimalla, P., Vasudevan, P., Carmeliet, P., Corbeil, D., Michalakis, S., Koch, E., et al. (2016). Stem Cell–Derived Photoreceptor Transplants Differentially Integrate Into Mouse Models of Cone-Rod Dystrophy. Invest. Ophthalmol. Vis. Sci. 57, 3509. 10.1167/iovs.16-19087.

10. Gagliardi, G., Ben M’Barek, K., Chaffiol, A., Slembrouck-Brec, A., Conart, J.-B., Nanteau, C., Rabesandratana, O., Sahel, J.-A., Duebel, J., Orieux, G., et al. (2018). Characterization and Transplantation of CD73-Positive Photoreceptors Isolated from Human iPSC-Derived Retinal Organoids. Stem Cell Reports 11, 665–680. 10.1016/j.stemcr.2018.07.005.

11. Carido, M., Völkner, M., Steinheuer, L.M., Wagner, F., Kurth, T., Dumler, N., Ulusoy, S., Wieneke, S., Norniella, A.V., Golfieri, C., et al. (2023). Reliability of human retina organoid generation from hiPSC-derived neuroepithelial cysts. Front. Cell. Neurosci. 17, 1166641. 10.3389/fncel.2023.1166641.

12. Tessmer, K., Borsch, O., Ader, M., and Gasparini, S.J. (2023). Micromanipulator-Assisted Subretinal Transplantation of Human Photoreceptor Reporter Cell Suspensions into Mice. In Brain Organoid Research Neuromethods., J. Gopalakrishnan, ed. (Springer US), pp. 81–98. 10.1007/978-1-0716-2720-4_5.

13. Zerti, D., Dorgau, B., Sernagor, E., Armstrong, L., Lako, M., and Hilgen, G. (2025). Evaluating the outcomes of pluripotent stem-cell-derived photoreceptor transplantation in retinal repair. The FEBS Journal, febs.70127. 10.1111/febs.70127.

14. Sullivan, L.S., and Daiger, S.P. RetNet. Retinal Information Network. https://retnet.org.

15. Ribeiro, J., Procyk, C.A., West, E.L., O’Hara-Wright, M., Martins, M.F., Khorasani, M.M., Hare, A., Basche, M., Fernando, M., Goh, D., et al. (2021). Restoration of visual function in advanced disease after transplantation of purified human pluripotent stem cell-derived cone photoreceptors. Cell Reports 35, 109022. 10.1016/j.celrep.2021.109022.

16. Lin, V., Lee, W., Kang, E.Y.-C., Liu, P.-K., and Wang, N.-K. (2026). Outer retinal tubulation associated with photoreceptor degeneration. Prog Retin Eye Res 111, 101435. 10.1016/j.preteyeres.2026.101435.

17. Sudharsan, R., Dolgova, N., Kwok, J., Gray, A., Sato, Y., Madrigal, A.L., Susaimanickam, P.J., Kriukov, E., Baranov, P., Wolfe, J.H., et al. (2025). Metabolic stress and early cell death in photoreceptor precursor cells following retinal transplantation. Stem Cell Res Ther 16, 397. 10.1186/s13287-025-04509-w.

18. Barber, A.C., Hippert, C., Duran, Y., West, E.L., Bainbridge, J.W.B., Warre-Cornish, K., Luhmann, U.F.O., Lakowski, J., Sowden, J.C., Ali, R.R., et al. (2013). Repair of the degenerate retina by photoreceptor transplantation. Proc. Natl. Acad. Sci. U.S.A. 110, 354–359. 10.1073/pnas.1212677110.

19. Kinouchi, R., Takeda, M., Yang, L., Wilhelmsson, U., Lundkvist, A., Pekny, M., and Chen, D.F. (2003). Robust neural integration from retinal transplants in mice deficient in GFAP and vimentin. Nat Neurosci 6, 863–868. 10.1038/nn1088.

20. Zhang, Y., Caffe, A.R., Azadi, S., Van Veen, T., Ehinger, B., and Perez, M.-T.R. (2003). Neuronal Integration in an Abutting-Retinas Culture System. Invest. Ophthalmol. Vis. Sci. 44, 4936. 10.1167/iovs.02-0640.

21. Matsuyama, T., Tu, H.-Y., Sun, J., Hashiguchi, T., Akiba, R., Sho, J., Fujii, M., Onishi, A., Takahashi, M., and Mandai, M. (2021). Genetically engineered stem cell-derived retinal grafts for improved retinal reconstruction after transplantation. iScience 24, 102866. 10.1016/j.isci.2021.102866.

22. Watanabe, M., Yamada, T., Koike, C., Takahashi, M., Tachibana, M., and Mandai, M. (2025). Transplantation of genome-edited retinal organoids restores some fundamental physiological functions coordinated with severely degenerated host retinas. Stem Cell Reports 20, 102393. 10.1016/j.stemcr.2024.102393.

23. Rich, K.A., Figueroa, S.L., Zhan, Y., and Blanks, J.C. (1995). Effects of müller cell disruption on mouse photoreceptor cell development. Experimental Eye Research 61, 235–248. 10.1016/S0014-4835(05)80043-0.

24. Wang, X., Iannaccone, A., and Jablonski, M.M. (2004). Contribution of Müller cells toward the regulation of photoreceptor outer segment assembly. Neuron Glia Biol. 1, 291–296. 10.1017/S1740925X05000049.

25. Völkner, M., Wagner, F., Steinheuer, L.M., Carido, M., Kurth, T., Yazbeck, A., Schor, J., Wieneke, S., Ebner, L.J.A., Del Toro Runzer, C., et al. (2022). HBEGF-TNF induce a complex outer retinal pathology with photoreceptor cell extrusion in human organoids. Nat Commun 13, 6183. 10.1038/s41467-022-33848-y.

26. Hammer, J., Röppenack, P., Yousuf, S., Schnabel, C., Weber, A., Zöller, D., Koch, E., Hans, S., and Brand, M. (2022). Visual Function is Gradually Restored During Retina Regeneration in Adult Zebrafish. Front. Cell Dev. Biol. 9, 831322. 10.3389/fcell.2021.831322.

27. Hanker, J.S., Deb, C., Wasserkrug, H.L., and Seligman, A.M. (1966). Staining Tissue for Light and Electron Microscopy by Bridging Metals with Multidentate Ligands. Science 152, 1631–1634. 10.1126/science.152.3729.1631.

28. Venable, J.H., and Coggeshall, R. (1965). A SIMPLIFIED LEAD CITRATE STAIN FOR USE IN ELECTRON MICROSCOPY. The Journal of Cell Biology 25, 407–408. 10.1083/jcb.25.2.407.

29. Hao, Y., Stuart, T., Kowalski, M.H., Choudhary, S., Hoffman, P., Hartman, A., Srivastava, A., Molla, G., Madad, S., Fernandez-Granda, C., et al. (2024). Dictionary learning for integrative, multimodal and scalable single-cell analysis. Nat Biotechnol 42, 293–304. 10.1038/s41587-023-01767-y.

