## Supplementary figures and images for "Transplanted human photoreceptors differentially survive, incorporate, and mature in mildly and severely degenerated mouse retinae"

### Supplemental Figure 1

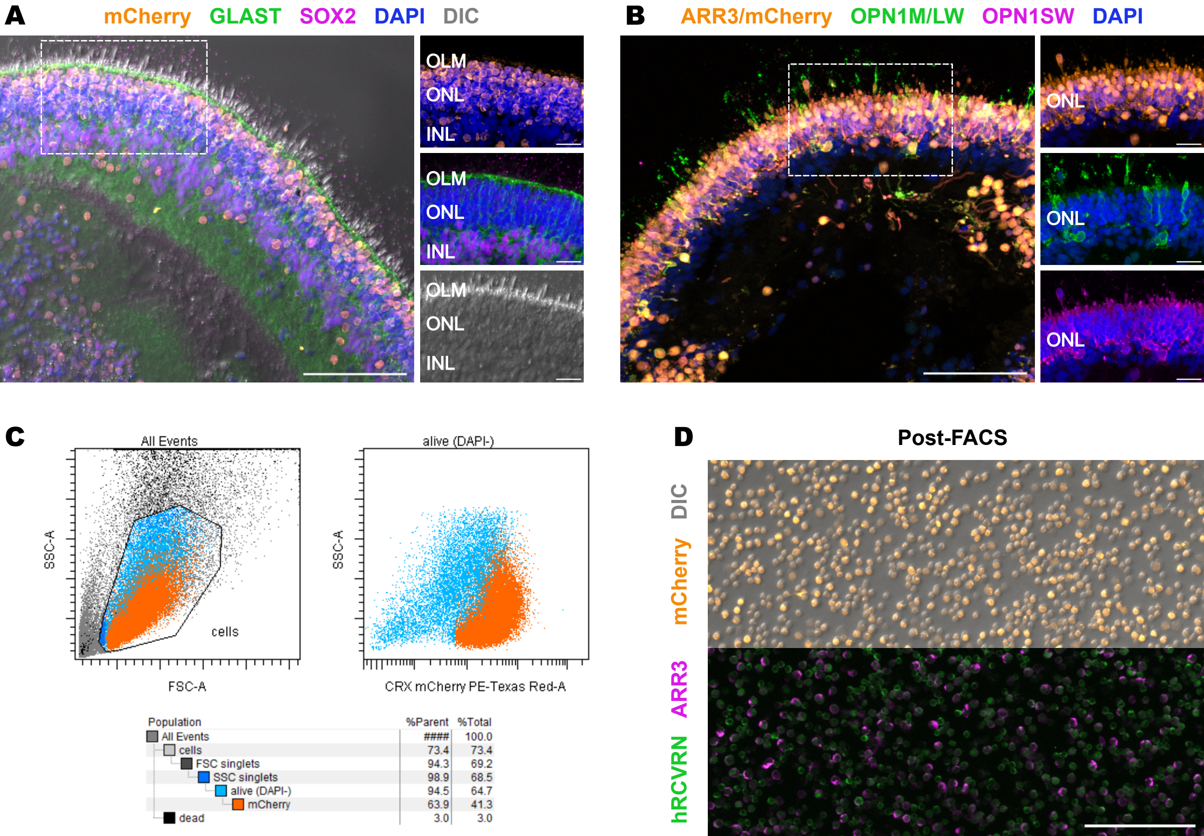

### Supplemental Figure 2

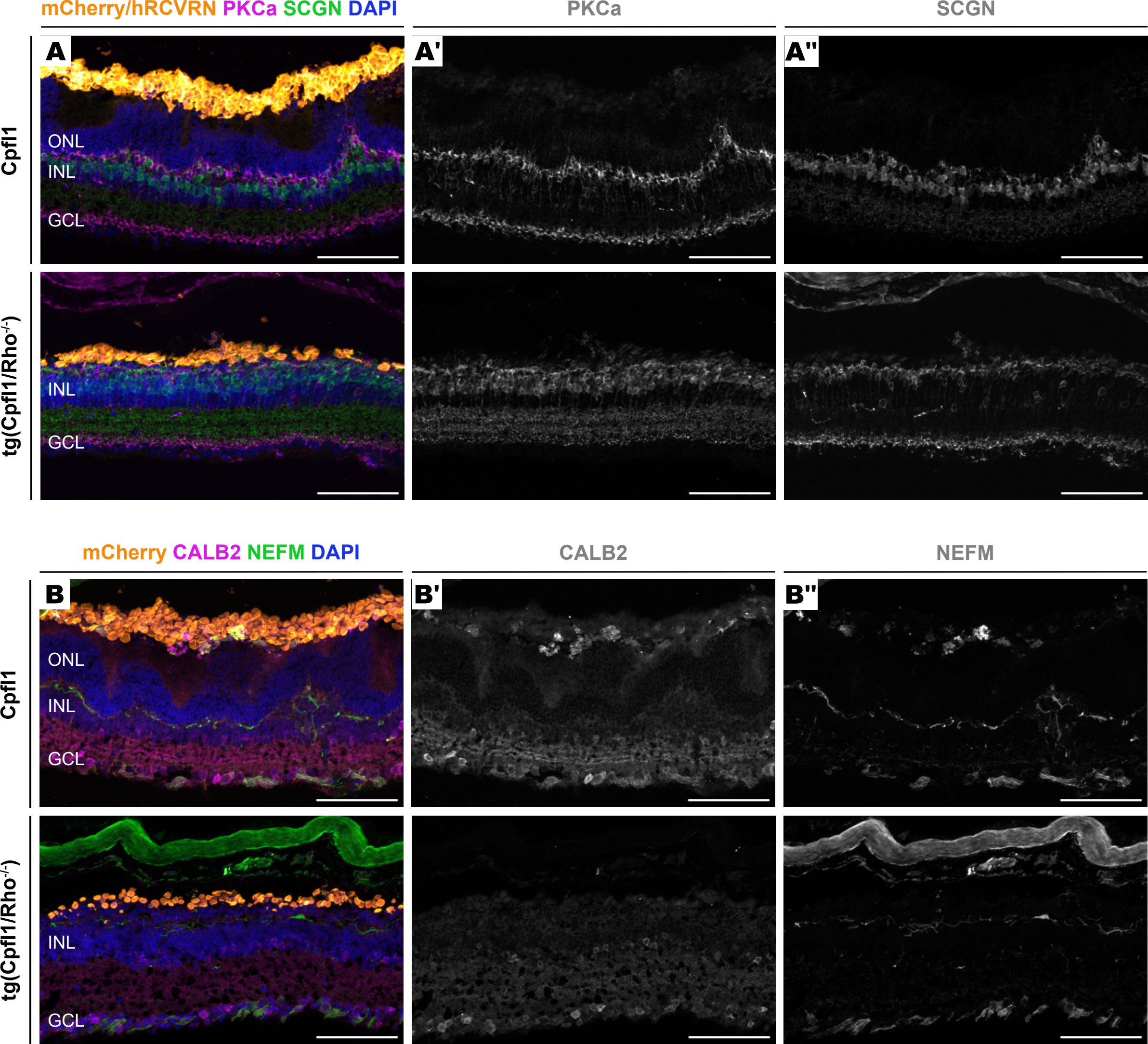

### Supplemental Figure 3

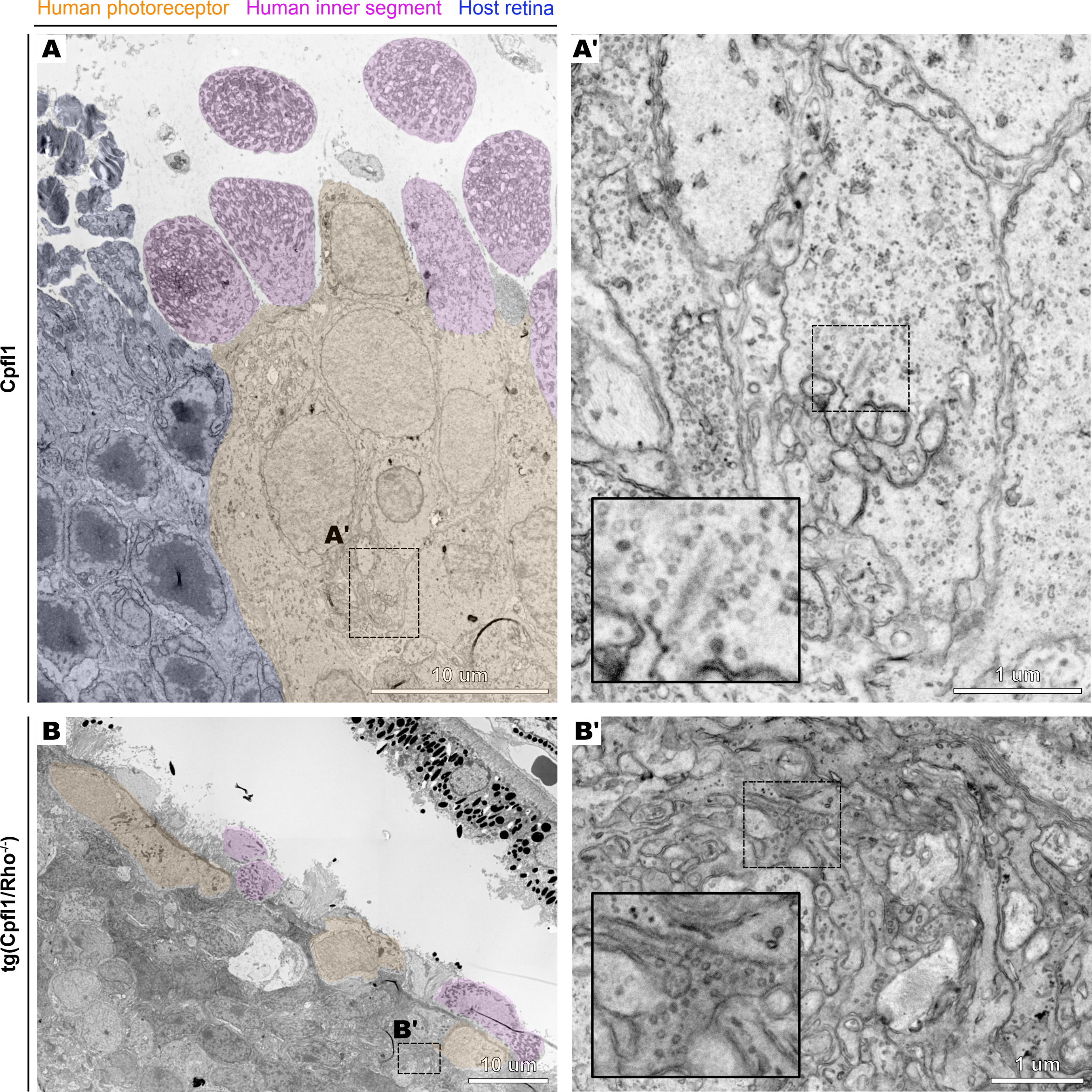

### Supplemental Figure 4

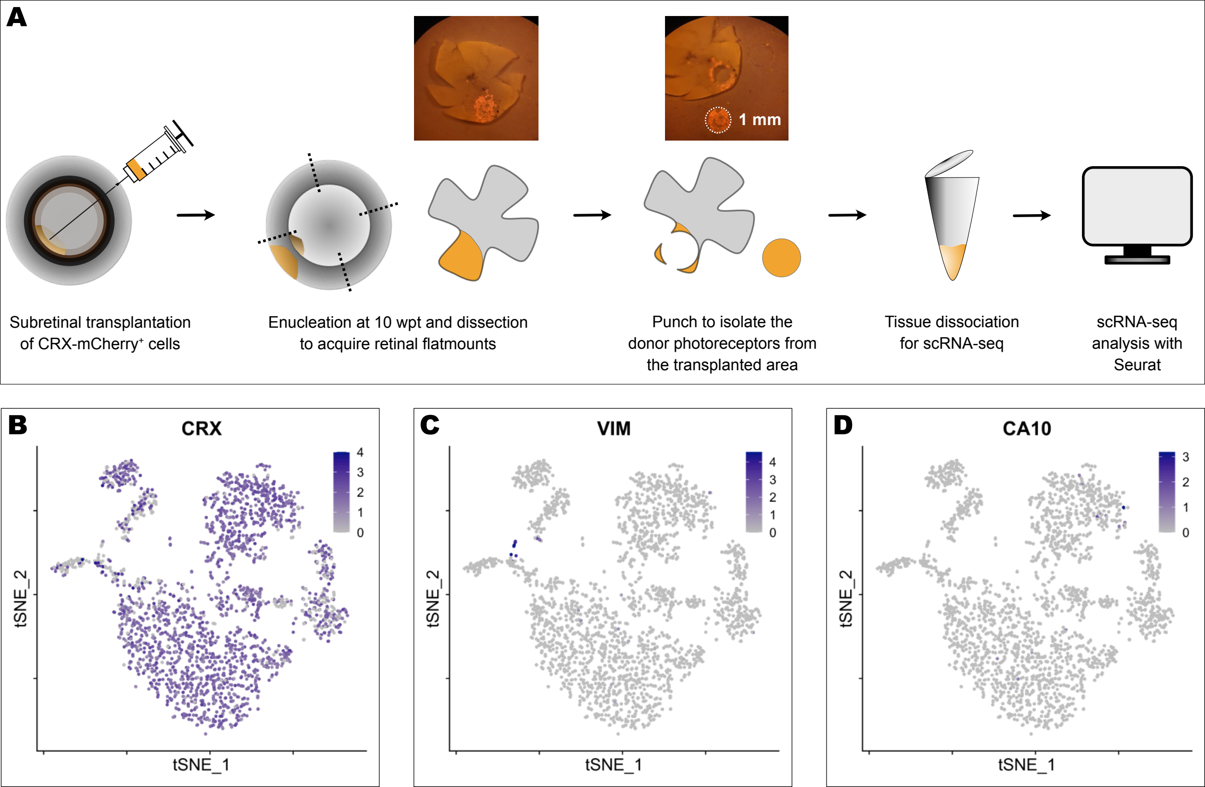

### Supplemental Figure 5

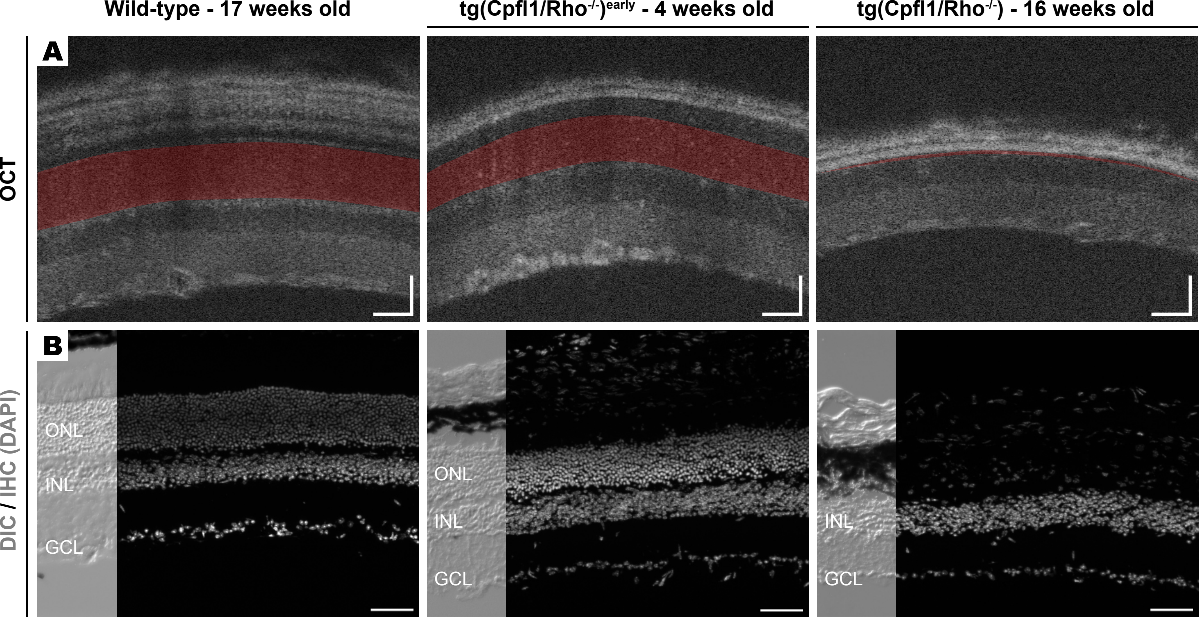
